# STID: A Standardized Spatial Transcriptomic Analysis Framework for Infectious Diseases

**DOI:** 10.64898/2026.05.24.727492

**Authors:** Yulong Qin, Yue Peng, Qi Chen, Jialing Chen, Peidi Ren, Haohao Deng, Daxi Wang, Xin Liu, Zhihua Ou, Ziqing Deng, Xiaofeng Shi

## Abstract

Spatial transcriptomic studies of infectious diseases still rely on fragmented data analysis processes. Here, we developed STID, a standardized framework for spatial transcriptomic analysis of infectious diseases that leverages the Seurat ecosystem and incorporates Python-based modules. STID provides an extensible infection-specific data structure and supports a full suite of analyses, such as pathogen background correction, infection-associated spot and niche identification, single-sample niche characterization, and multi-sample comparative and temporal analyses. Moreover, STID is broadly applicable to spatial transcriptomic data from infectious diseases caused by bacteria, viruses, and parasites, and enables systematic characterization of the structural features, cellular composition, molecular functions, and host–pathogen interactions within pathogen-infected and/or host-responsive niches. Overall, STID provides an accessible, reproducible, and extensible framework for analyzing infection-associated spatial transcriptomic data and for dissecting host–pathogen interactions in their native spatial microenvironments.

**Motivation:** Spatial transcriptomics technologies have emerged as powerful approaches for dissecting the structural and functional features of spatial microenvironments. However, the current general-purpose tools remain fundamentally inadequate for resolving the spatial heterogeneity of infectious disease samples, where the intricacies of host–pathogen interactions render spatial microenvironments both challenging to dissect and largely inaccessible. Tools tailored to infectious diseases are critically lacking, including those for reducing pathogen-derived background noise, identifying and isolating infection□associated spots or niches, dissecting host–pathogen interactions, and supporting systematic multi-sample analyses. We therefore developed STID, a unified framework that integrates standardized workflows and addresses the analytical bottlenecks in spatial transcriptomic analysis of infectious diseases.

**Highlights:**

- STID standardizes spatial transcriptomic analysis in infectious diseases
- STID improves pathogen-infected spot detection by correcting pathogen background
- STID distinguishes pathogen-infected and host-responsive niches
- STID supports multi-sample comparative and temporal analyses of niches

**Graphical abstract:** 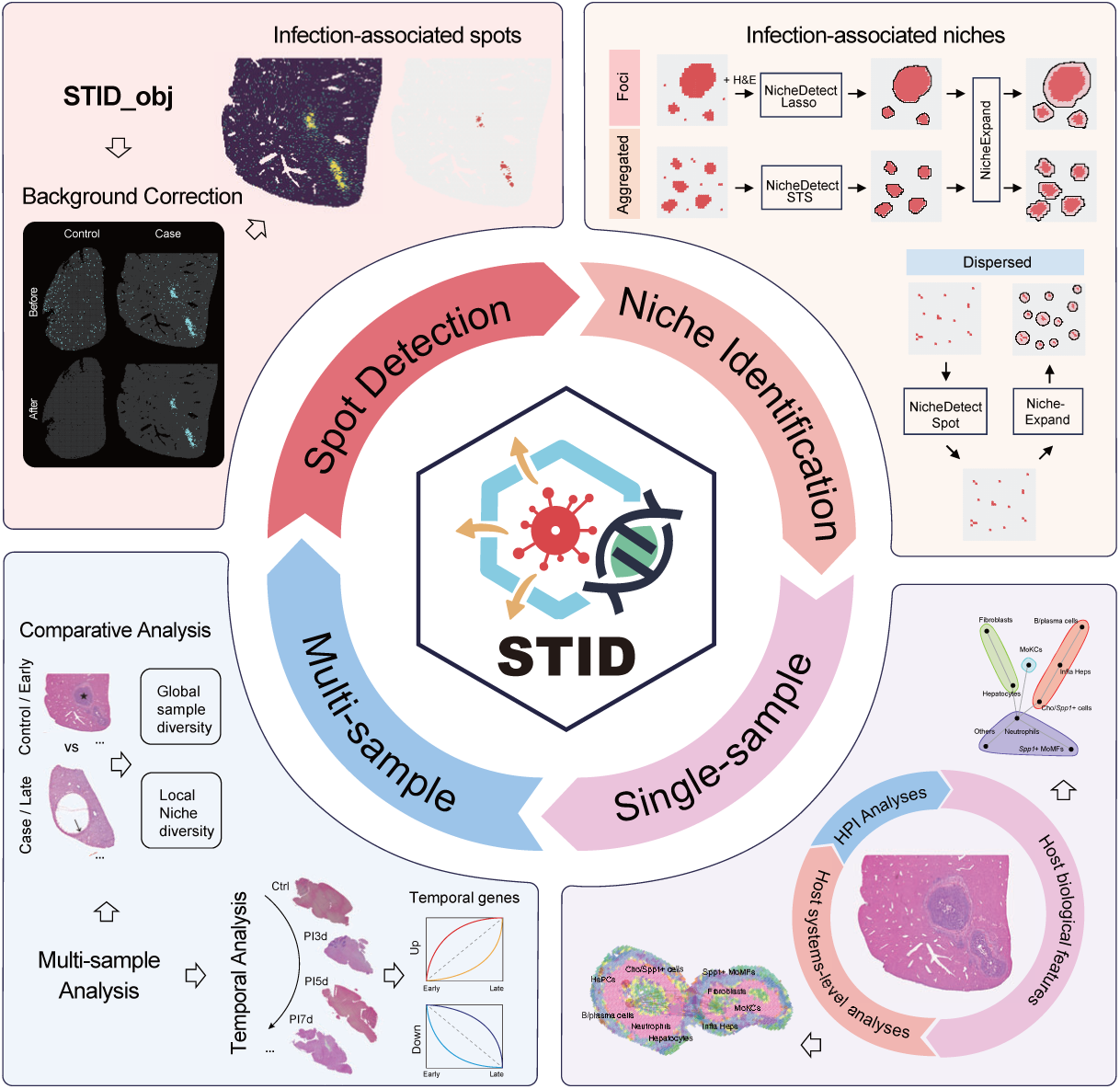

## Introduction

Spatial biology provides a new dimension for understanding the complex organization of living systems through characterization of the in situ distribution of cells and molecules^1,2^. Spatial transcriptomics technologies enable quantitative transcriptome-wide profiling while preserving tissue spatial information^3,4^ and have been widely applied in developmental biology^5^, cancer research^6^, and neuroanatomy^7^, thereby facilitating systematic analysis of the structural and functional features of spatial microenvironments. In infectious diseases, host–pathogen interactions are highly dependent on local spatial niches. Pathogen colonization and dissemination, immune cell recruitment and activation, and tissue injury and repair exhibit pronounced spatial heterogeneity and temporal dynamics^4,8–14^. Consequently, spatial transcriptomics technologies have been widely applied in infectious disease research, providing high-resolution approaches for characterizing the structural and functional features of spatial microenvironments, elucidating disease mechanisms, and identifying potential therapeutic targets^15–17^.

Despite the broad applicability of spatial transcriptomics, specialized analytical frameworks for infectious disease research remain scarce. The current general-purpose tools, such as Seurat^18^, Scanpy^19^, Squidpy^20^, and Stereopy^21^, lack infection-specific modules and provide inadequate support for reducing pathogen-derived background noise, detecting infection-associated spots and identifying and isolating infection-associated spatial niches. Moreover, existing tools poorly support multi-sample comparative and temporal analyses in infectious diseases. Consequently, researchers have to rely on fragmented cross-platform workflows, which inevitably increase analytical complexity and undermine reproducibility.

To address these limitations, we developed STID (Spatial Transcriptomics toolkit for Infectious Diseases), a standardized spatial transcriptomic analysis framework specifically designed for infectious diseases. STID leverages the Seurat ecosystem and incorporates Python-based modules. Moreover, STID enables pathogen background correction, detection of infection-associated spots, and identification of infection-associated spatial niches. In this study, we define infection-associated spatial niches as spatially organized tissue regions composed of pathogen-infected spots, host-responsive spots, or both. At the single-sample level, STID enables researchers to characterize the infection-associated niches across spatial organization, cellular composition, intercellular communication, and gene regulatory networks. In addition, STID supports multi-sample comparative analysis and temporal analysis, thereby systematically depicting the spatiotemporal dynamics of host–pathogen interactions.

Unlike general-purpose tools, STID is tailored to infectious disease research, offering a full workflow that spans from identification and characterization of infection-associated niches in single-sample to comparative and temporal analyses in multi□sample (Table 1). In this study, we applied STID to publicly available spatial transcriptomic datasets from infectious diseases, including alveolar echinococcosis (AE)^12^ and Japanese encephalitis (JE)^9^. The results establish STID as a powerful framework for spatial transcriptomic analysis of infectious diseases. Specifically, STID enables robust characterization of infection-associated niches, investigates host–pathogen interaction mechanisms, and prioritizes potential therapeutic targets, thereby validating its broad utility and analytical capabilities. Overall, STID serves as a convenient, efficient, and reproducible framework for spatial transcriptomic analysis of infectious diseases, providing computational support for pathogen biology, immunology, and translational medicine research.

## Results

### STID establishes a unified framework for spatial transcriptomics of infectious diseases

STID integrates input/output, preprocessing, and downstream analysis modules, establishing an end-to-end workflow from raw spatial transcriptomic data input to infection-associated single-sample and multi-sample analyses (Figure 1A). Spatial transcriptomic data are broadly classified into square-grid and hex-grid spatial layouts^20^. Representative square-grid platforms include Stereo-seq, Visium HD, and Slide-seq, whereas standard Visium represents a typical hex-grid platform. STID accepts both square-grid and hex-grid spatial transcriptomic data, and supports bidirectional conversion between H5AD and Seurat RDS formats. During preprocessing, STID integrates cell type annotation and data normalization. The Seurat object is then transformed into an infection-specific STID class, thereby providing a standardized foundation for downstream infection-associated analyses.

**Figure 1.**
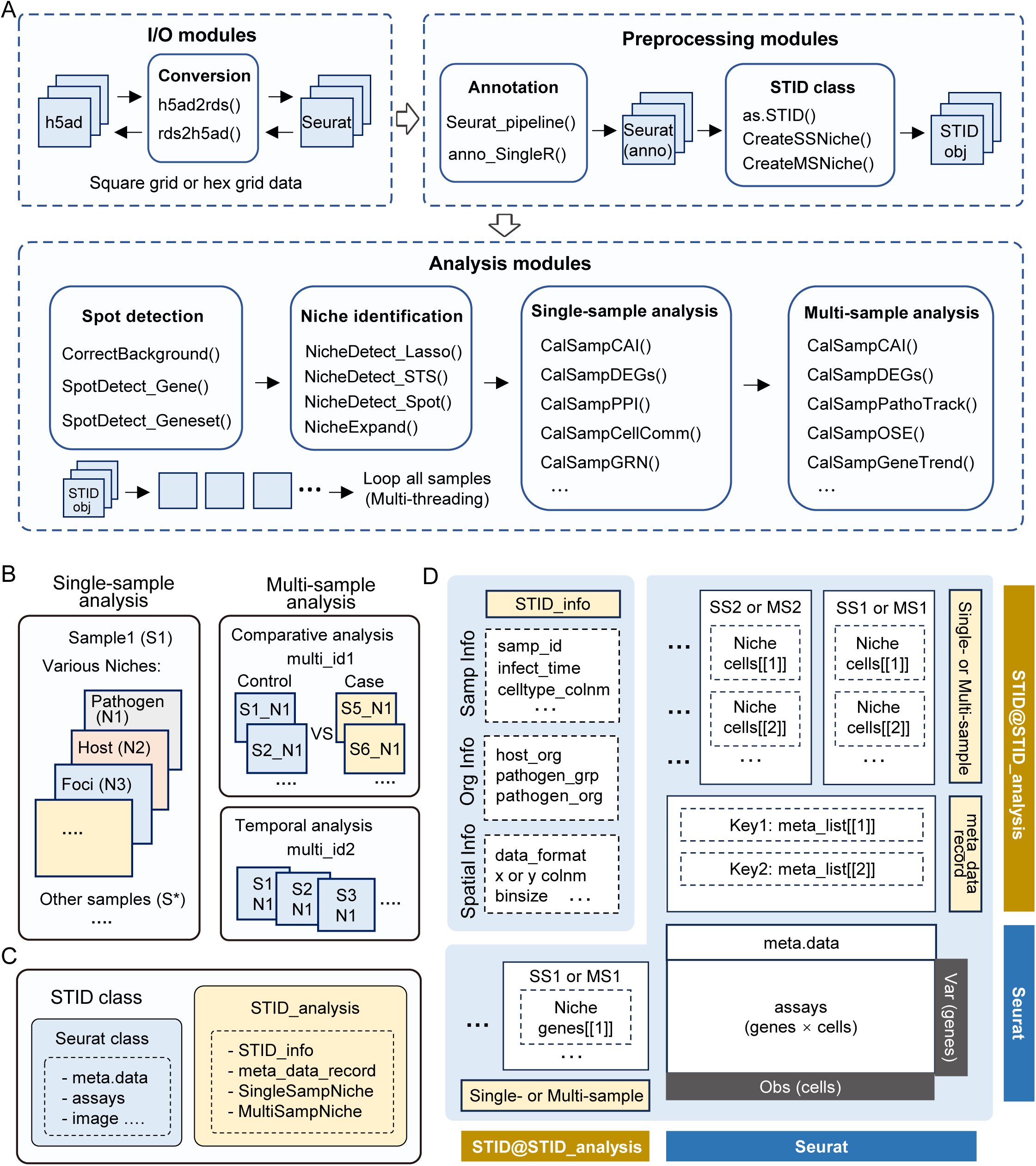
Analysis workflow and data structure design of STID. (A) Overall workflow of STID. The workflow comprises input/output, preprocessing, and analysis modules. The input/output modules convert square-grid and hex-grid spatial transcriptomic data between H5AD and Seurat RDS formats. Preprocessing includes cell type annotation and conversion of Seurat data to the STID format. Analysis modules perform infection-associated spot detection, infection-associated niche identification, and single- and multi-sample analyses. (B) Framework for single- and multi-sample analyses in STID. Single-sample analysis characterizes spatial organization patterns of different types of infection-associated niches. Multi-sample analysis includes comparative and temporal strategies to assess differences across groups and spatiotemporal dynamics during infection. (C) Design of the STID class. The STID class is built on the Seurat class and incorporates a new STID analysis slot to store infection-associated spatial transcriptomics analysis results. This slot contains four modules: STID_info, meta_data_record, SingleSampleNiche, and MultiSampNiche. (D) Composition of the STID class. STID_info records sample, host–pathogen, and spatial transcriptomic platform information. The meta_data_record module stores analysis-derived metadata and supports bidirectional conversion with Seurat meta.data. SingleSampleNiche and MultiSampNiche store cellular- and gene-level results obtained from single-sample and multi-sample niche analyses.

For downstream analysis, STID provides four core functional modules that address key questions in infectious diseases, including pathogen background correction, identification of infection-associated spots and niches, single-sample analysis, and multi-sample analysis. Single-sample analysis characterizes structural and functional features of infection-associated niches (pathogen-infected niches and/or host-responsive niches), whereas multi-sample analysis performs comparative and temporal analyses to systematically characterize the group differences and reveal spatiotemporal niche dynamics during infection progression (Figures 1A and 1B). Furthermore, STID uses parallel computing to accelerate key analytical steps.

To support the above analytical framework, STID defines a unified data structure to store and organize infection-associated spatial transcriptomic results (Figure 1C). The STID class extends the Seurat class and adds a new analysis slot. This slot comprises four modules: (1) STID_info, which records sample, host–pathogen, and spatial transcriptomic platform information; (2) meta_data_record, which stores analysis-derived metadata and supports bidirectional conversion with Seurat meta.data; (3) SingleSampleNiche and (4) MultiSampNiche, which store cell- and gene-level results for single-sample and multi-sample niches, respectively (Figure 1D).

### STID corrects pathogen-derived background noise and improves pathogen-infected spot detection

To address pathogen-derived background noise in spatial transcriptomic data of infectious diseases^22,23^, STID introduces a background correction strategy based on constraining the distribution of pathogen gene expression in control samples, thereby improving the specificity and robustness of pathogen-infected spot detection. In the AE model, pathogen gene expression was widely dispersed in control samples before correction, indicating substantial pathogen background noise. After pathogen background correction, nonspecific signals were effectively eliminated in control samples, whereas the spatial aggregation patterns were preserved and further enhanced in infected samples (Figure 2A). This correction also reduced the distribution of abnormally high pathogen gene expression in control samples (Figure 2B), as well as both the overall pathogen gene expression and the number of positive spots (Figures 2C and 2D). In infected samples, these metrics remained largely unchanged. Collectively, these results indicate that STID reduces background noise while preserving authentic infection-derived signals. Furthermore, in the tuberculosis (TB) model^4^, this background correction strategy showed consistent and robust performance (Figures S1A–S1D), further supporting the general applicability of STID across pathogen types and tissue infection contexts.

**Figure 2.**
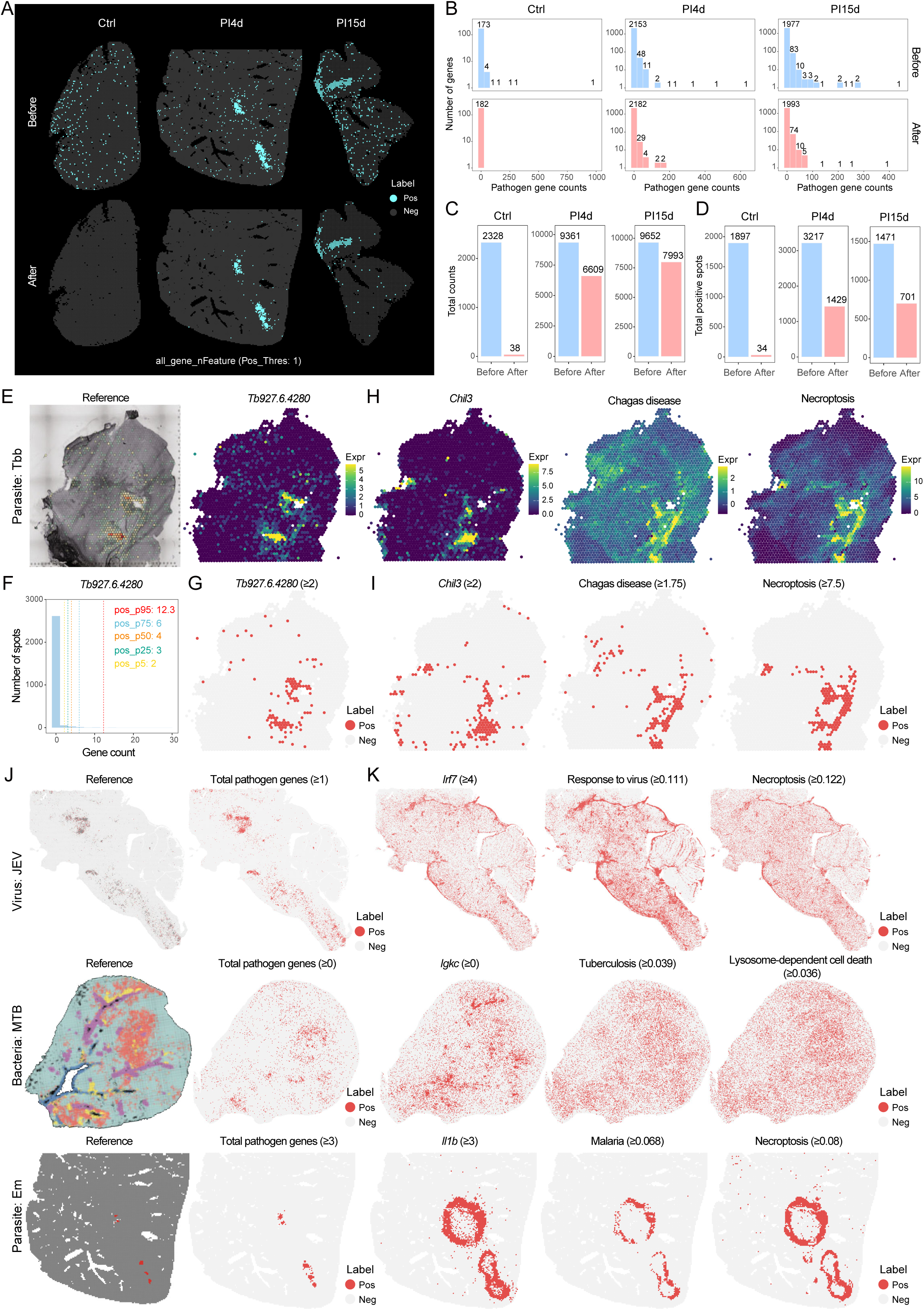
Pathogen background correction and infection-associated spot detection. (A) Spatial maps showing pathogen gene expression in AE model samples before and after pathogen background correction (Control, PI4d, and PI79d stages). Top: before correction; bottom: after correction.(B) Histograms showing frequency distributions of total pathogen gene counts in samples at different infection stages. Top: before correction; bottom: after correction. (C–D) Bar plots showing changes in total pathogen gene counts and numbers of pathogen-infected spots across infection stages before and after pathogen background correction. (E) Spatial maps showing the distribution of *Trypanosoma brucei Tb927.6.4280* expression. Left: published study; right: STID analysis. (F) Histograms showing the distribution of spot numbers across *Tb927.6.4280* count levels. (G) Spatial maps showing *Tb927.6.4280*-positive spots defined by expression threshold. (H) Spatial maps showing host response features, including *Chil3* expression, Chagas disease-related signature scores, and necroptosis gene set scores. (I) Spatial maps showing the distribution of host-responsive spots identified using *Chil3* expression, Chagas disease scores, or necroptosis scores. (J) Spatial maps showing the distribution of pathogen-infected spots across infection models (Japanese encephalitis, tuberculosis, and alveolar echinococcosis). Left: published studies; right: STID analysis. (K) Spatial maps showing the distribution of host-responsive spots across infection models, identified using representative host gene expression, anti-pathogen response scores, and cell death scores.

To detect infection-associated spots, we used STID to profile the spatial distribution of a representative pathogen gene (*Tb927.6.4280*) in the African trypanosomiasis model^14^ (Figure 2E), and defined the infection threshold using the expression intensity distribution of this gene (Figure 2F). Positive spots identified by this threshold exhibited pronounced spatial clustering and showed strong concordance with known pathogen-infected spots (Figure 2G), demonstrating that STID robustly reconstructs infection-associated spots. To dissect the spatial relationship between pathogen-infected spots and host-responsive spots, we further analyzed the spatial distribution of host response features. *Chil3* expression, Chagas disease-related signature scores, and necroptosis scores all exhibited locally enriched spatial patterns (Figure 2H), enabling the identification of host-responsive spots. Notably, these host-responsive spots showed strong spatial overlap with the pathogen-infected spots (Figure 2I).

Next, we identified the pathogen-infected spots based on the respective pathogen gene expression thresholds in the JE, TB, and AE models. Our results showed high concordance with the published study results, confirming STID’s robustness across pathogen types (Figure 2J). Further analysis revealed that host-responsive spots only partially overlapped with the pathogen-infected spots across infection conditions. The strongest host responses did not necessarily coincide with the highest pathogen burden (Figure 2K), indicating pronounced spatial compartmentalization between pathogen-infected spots and host-responsive spots. For spatial transcriptomic datasets lacking pathogen information, including viral myocarditis (VMC)^11^, Venezuelan equine encephalitis (VEE)^10^, *Klebsiella pneumoniae* pneumonia^8^, and malaria^13^, host-responsive spots were readily identified based on the spatial distribution of host response features (Figures S1E and S1F), demonstrating the broad applicability of STID across diverse spatial transcriptomic datasets.

### STID identifies infection-associated niches with distinct spatial patterns

Different pathogen types and tissue/organ contexts contribute to the pronounced spatial heterogeneity of infectious diseases^4,8,9,11,12,14^. To systematically characterize infection-associated niches with distinct spatial distribution patterns, STID establishes three niche identification modes based on the spatial aggregation features of infection-associated spots: Foci, Aggregated, and Dispersed (Figure 3A). The Foci mode provides the interactive delineation of infection-associated niches through the NicheDetect_Lasso Shiny application based on hematoxylin and eosin (H&E)-stained histological information, pathogen-infected spots, and host-responsive spots (Figures 3B and S2A). The Aggregated mode enables automatic identification of niches with clustered distributions of positive spots using NicheDetect_STS (Figure S2B), whereas the Dispersed mode directly defines all positive spots as niches. All three modes support flexible expansion of niche boundaries through NicheExpand (Figure 3A).

**Figure 3.**
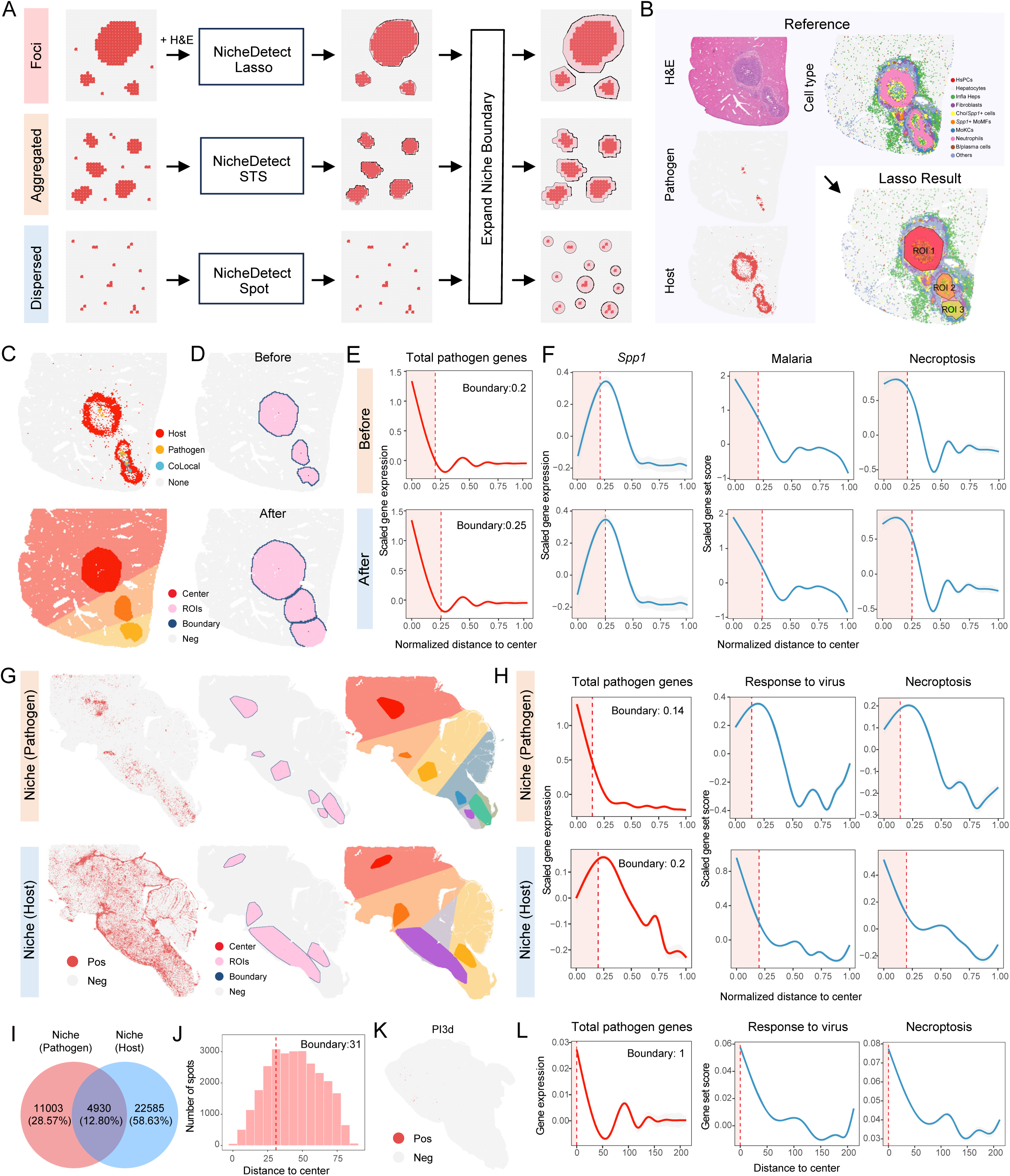
Identification of infection-associated niches. (A) Schematic overview of niche identification strategies, including Foci, Aggregated, and Dispersed types. Foci-type niches are identified using NicheDetect_Lasso incorporating H&E histological information; Aggregated-type niches are identified using NicheDetect_STS; Dispersed-type niches are identified using NicheDetect_Spot. All identified niches can be further expanded using NicheExpand. (B) Workflow of the NicheDetect_Lasso method. Gold-standard H&E information is integrated with multiple auxiliary features, including pathogen-infected spots, host-responsive spots, and cell type annotations, to accurately identify infection-associated niches. (C) Spatial maps showing the spatial relationship between pathogen-infected spots, host-responsive spots, and identified niches in the AE model. Top: colocalization of pathogen-infected and host-responsive spots. Bottom: multiple ROIs within the niche and corresponding bystander regions. Dark areas indicate niche regions and light areas indicate bystander regions. (D) Spatial maps showing niche structures before and after boundary expansion. Red indicates center points and blue indicates boundaries. Top: before expansion; bottom: after expansion. (E) Fitted curves showing spatial gradient of total pathogen gene counts from niche centers toward the periphery before and after boundary expansion. Dashed lines indicate niche boundaries. (F) Fitted curves showing spatial gradient of host response features (*Spp1* expression, malaria-related signature scores, and necroptosis scores) from niche centers toward the periphery. (G) Spatial maps showing pathogen-infected niche and host-responsive niche in JE model samples. Top: pathogen-infected niche; bottom: host-responsive niche. Left: positive spot distribution; middle: niche structures; right: multiple ROIs within the niche and corresponding bystander regions. (H) Fitted curves showing spatial gradient of total pathogen gene counts, response to virus, and necroptosis from niche centers toward the periphery. Dashed lines indicate niche boundaries. (I) Venn diagram showing spatial overlap between pathogen-infected and host-responsive niches. (J) Histogram showing the distribution of distances from host-responsive niche spots to the nearest pathogen-infected niche center. Dashed line indicates the boundary of the pathogen-infected niche. (K) Spatial maps showing the distribution of pathogen-infected spots. (L) Fitted curves showing spatial gradient of total pathogen gene counts, response to virus, and necroptosis from niche centers toward the periphery. Dashed lines indicate niche boundaries.

In the AE model, we applied the Foci mode of STID for niche identification followed by boundary expansion. The niche prior to expansion exhibited a spatial pattern largely consistent with the union of pathogen-infected and host-responsive spots (Figure 3C), and delineated the center and boundary of different regions of interest (ROIs) within the niche (Figures 3C and 3D). Spatial gradient analysis of gene expression and functional scores revealed that pathogen gene expression gradually decreased from center to boundary in the niche, reaching the lowest levels near the expanded niche boundary (Figure 3E), suggesting that niche boundaries can be finely adjusted based on pathogen gene expression gradients. In contrast, host response features exhibited an opposing spatial gradient, with peak signals localized near the niche boundary (Figure 3F). Our results reveal the distinct spatial organization between pathogen-infected and host-responsive niches.

In the JE model, we applied the Aggregated mode of STID for niche identification followed by boundary expansion. After spatial low-density filtering of positive spots, infection-associated and host-responsive niches were identified (Figures 3G and S2C). Spatial gradient analysis of gene expression and functional scores revealed that pathogen gene expression peaked near the niche center, whereas host response features peaked near the niche boundary within pathogen-infected niches (Figure 3H). After boundary expansion, the niche encompassed a more complete region of high pathogen expression (Figures S2D and S2E). In host-responsive niches, an opposite pattern was observed, with pathogen gene expression peaking near the niche boundary and host response features peaking near the niche center. This spatial organization was further validated in TB and African trypanosomiasis models (Figures S2F, S2G, S2J, and S2K). Additionally, pathogen-infected and host-responsive niches showed low spatial overlap, and host-responsive niches exhibited broader spatial coverage. In different infection models, we observed that lower overlap between pathogen-infected and host-responsive niches was associated with greater divergence in the spatial gradient patterns of pathogen gene expression and host-response-related functional scores between pathogen-infected and host-responsive niches (Figures 3I, S2H, and S2L). Distance-based analysis further showed that host-responsive niches were primarily distributed near the boundaries of pathogen-infected niches rather than regions of peak pathogen gene expression (Figures 3J, S2I, and S2M).

In addition, we applied the Dispersed mode of STID to define all pathogen-infected spots in the PI3d samples of the JE model as a niche. Spatial gradient analysis revealed concordant spatial gradients of pathogen gene expression and host response features (Figures 3K and 3L). For spatial transcriptomic datasets lacking pathogen information, such as *Klebsiella pneumoniae* pneumonia and VMC, STID also enabled identification of host-responsive niches based on host response gene expression and functional scores (Figures S2N and S2O). The host response features exhibited an expected spatial gradient that gradually decreased from center to boundary.

### STID enables multiscale systems characterization of infection-associated niches

Characterizing the structural and functional features of infection-associated niches is essential for elucidating structure–function relationships. To this end, STID provides a niche-based systems analysis framework that enables a multiscale dissection of these niches across host biological features (structural, cellular, and molecular features), host systems-level organization (cell–cell communication and gene regulatory networks), and host–pathogen interactions (Figure 4A). In the AE model, the infection-associated niches were predominantly localized in the hepatic central vein region, whereas liver zonation in bystander regions remained relatively homogeneous (Figure 4B), suggesting specific tissue tropism of the pathogen. Cellular composition analysis further demonstrated that inflammation-associated cell types (e.g., neutrophils, *Spp1*□ MoMFs, MoKCs, and fibroblasts) were significantly enriched in infection-associated niches (Figure 4B). Cell aggregation index and cell high-aggregation region proportion further confirmed these enrichments (Figures 4C and 4D), indicating the formation of a highly organized inflammatory structure within the niche. Spatial colocalization analysis revealed a significant spatial proximity between neutrophils and *Spp1*□ MoMFs, with *Spp1*□ MoMFs predominantly localized around neutrophils (Figures 4E, S3A, and S3B). These results indicate that cells within the niche were not randomly distributed but instead exhibited a hierarchical spatial organization. Spatial gradient analysis further showed that neutrophils were predominantly distributed in the center of the niche, whereas *Spp1*□ MoMFs were predominantly distributed near the boundary (Figure 4F). This organization suggests distinct functional roles for different cellular populations within infection-associated niches.

**Figure 4.**
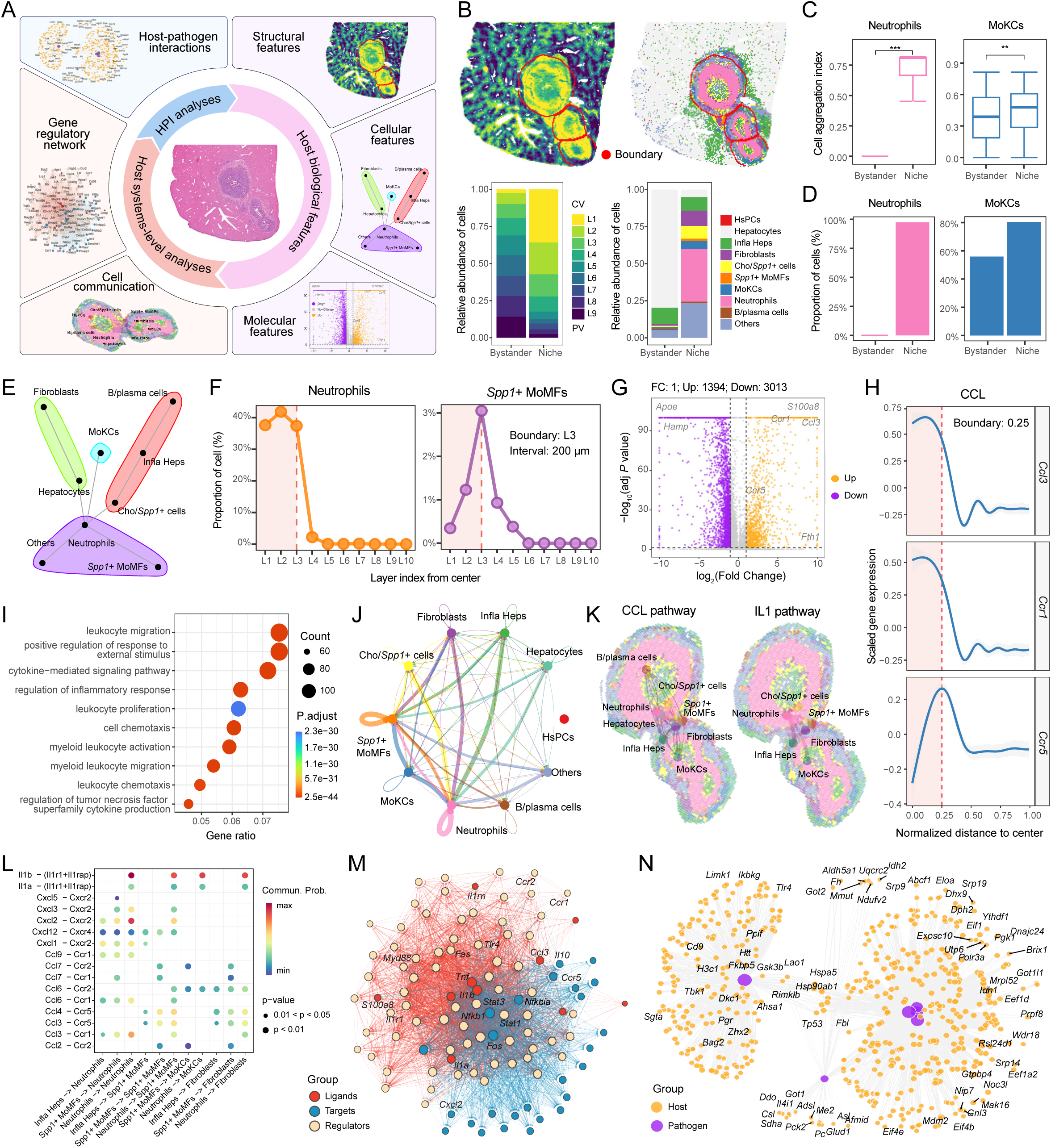
Multiscale single-sample niche analysis reveals host–pathogen interaction mechanisms. (A) Schematic overview of the single-sample niche analysis framework, comprising three major modules: Host biological features, Host systems-level analyses, and Host-pathogen interaction (HPI) analyses. Host biological features include structural features, cellular features, and molecular features. Host systems-level analyses include cell–cell communication analysis and gene regulatory network inference. (B) Spatial maps and stacked bar plots showing liver zonation and cellular composition in the AE model. Left: liver zonation; right: cellular composition. Top: spatial maps, with red circles indicating niche boundaries; bottom: stacked bar plots. (C) Boxplots showing the cell aggregation index in the niche and bystander region. Left: neutrophils; right: MoKCs. (D) Bar plots showing the proportion of cells located in high-aggregation regions within the niche and bystander region. Left: neutrophils; right: MoKCs. (E) Network diagram showing cell–cell interaction networks within the niche inferred using mistyR, with cells within the same colored circle belonging to the same community module. (F) Line plots showing spatial gradient of cell type proportions from niche centers toward the periphery. Dashed lines indicate niche boundaries. Left: *Spp1*□ MoMFs; right: neutrophils. (G) Volcano plot showing differentially expressed genes between the niche and bystander region. (H) Fitted curves showing spatial gradients of differentially expressed genes *Ccl3*, *Ccr1*, and *Ccr5* from niche centers toward the periphery. Dashed lines indicate niche boundaries. (I) Bubble plot showing GO terms enriched among upregulated differentially expressed genes between the niche and bystander region. (J) Network diagram showing cell–cell communication weights among all cell types within the niche. (K) Spatial network maps showing cell–cell communication weights in CCL and IL1 signaling pathways within the niche. Left: CCL pathway; right: IL1 pathway. (L) Bubble plot showing ligand–receptor interaction strengths between cell types within the niche. (M) Regulatory network showing inferred ligand–regulator–target gene relationships between sender and target cells. Sender cells include inflammatory hepatocytes, neutrophils, and *Spp1*□ MoMFs; target cells include neutrophils, *Spp1*□ MoMFs, MoKCs, and fibroblasts. Red nodes: ligand genes; yellow nodes: regulators; blue nodes: target genes. (N) Protein–protein interaction network showing inferred host–pathogen interactions. Purple nodes: pathogen proteins; yellow nodes: host proteins.

Along with the establishment of spatial organization, the niche also underwent pronounced molecular and functional remodeling. Inflammatory genes (e.g., *S100a8*, *Ccl3*, *Ccr1*, and *Ccr5*) were significantly upregulated in the niche relative to bystander regions (Figures 4G and S3C), indicating a transition from homeostasis to inflammatory activation. Moreover, the upregulation of *Hamp* in bystander regions and upregulation of *Fth1* in the niche suggested that hepatic cells enhance iron storage and restrict iron release in the niche to counteract *Echinococcus multilocularis* invasion through a *Hamp*-mediated pathway. Gene Ontology (GO) enrichment analysis further revealed that the upregulated genes in the niche were primarily involved in immune-related processes, including leukocyte migration and myeloid leukocyte activation (Figure 4I). Moreover, the expression of *Ccl3* and *Ccr1* peaked near the niche center, whereas *Ccr5* expression was enriched near the niche boundary (Figures 4H and S3D). Consistently, myeloid leukocyte activation scores peaked near the niche center, whereas myeloid leukocyte migration scores were enriched near the boundary (Figure S3E). These findings indicate coordinated spatial layering of cellular, molecular, and functional features within niches, characterized by immune cell recruitment at the boundary and activation in the niche center.

Cell–cell communication analysis revealed interactions underlying the hierarchical spatial organization of niches (Figure 4J). CCL signaling was broadly distributed across various cell types, whereas IL1 signaling was predominantly produced by neutrophils and targeted other cell populations (Figure 4K), suggesting that neutrophils function as key signaling hubs within infection-associated niches. Further ligand–receptor analysis demonstrated that neutrophils coordinated inflammatory cell recruitment through the Il1a/Il1b–(Il1r1+Il1rap), Cxcl2–Cxcr2, and Ccl3/Ccl4–Ccr5 signaling axes, while also activating *Spp1*□ MoMFs and fibroblasts for tissue repair and remodeling (Figures 4L, S3F, and S3G). Based on these intercellular signaling networks, transcriptional regulatory programs were constructed and found to be driven by inflammatory ligands (e.g., *Il1a/Il1b* and *Ccl3*), mediated through key regulators (e.g., NF-κB, STAT*, Il1r1*, and *Ccr1*), and targeted downstream genes (Figures 4M and S3H). These findings suggest a continuous regulatory axis linking cell–cell communication to transcriptional responses within infection-associated niches.

Pathogens can navigate complex host environments to thrive, replicate, and ultimately transmit to new hosts. Pathogen–host gene correlation analysis showed that the expression of most host genes was negatively correlated with the expression of pathogen genes (Figure S3I), suggesting a substantial perturbation of host gene expression within infection-associated niches. By contrast, positively correlated genes were mainly enriched in lipid metabolism–related processes (Figure S3J), suggesting that *Echinococcus multilocularis* supports its survival by remodeling the host metabolic environment. This pattern is also consistent with the predominant localization of *Echinococcus multilocularis* in the central vein region of the liver (Figure 4B). Furthermore, protein–protein interaction analysis suggested that pathogen-associated proteins affect host ribosome biogenesis and protein translation, thereby playing a key role in pathogen replication and immune evasion (Figures 4N and S3K).

Here, STID established a systematic framework to characterize the infection-associated niches in the AE model, from tissue architecture to molecular mechanisms. Unlike the published study^12^, STID uses the infection-associated niche as the central analytical unit and provides more detailed evidence, thereby improving our understanding of host–pathogen interaction mechanisms during *Echinococcus multilocularis* infection.

### STID reveals spatial segregation of pathogen-infected and host-responsive niches

Next, we applied STID to systematically characterize the structural and functional differences between pathogen-infected and host-responsive niches in the JE model. The two niches showed distinct spatial distributions (Figure S4A). Compared with bystander regions, the pathogen-infected niche showed significantly increased proportions of the thalamus (TH) and hindbrain (HB) (Figures S4B, S4C). Spatial colocalization analysis revealed that NK and T cells exhibited stronger spatial proximity within pathogen-infected niches (Figure S4D). Moreover, pathogen-infected niches showed marked upregulation of viral genes (e.g., *NS1* and *NS2a*) (Figure S4F), and the upregulated genes were enriched in neural function–related pathways (Figure S4G). Overall, the pathogen-infected niches are pathogen-resident regions characterized by local immune activation.

Compared with bystander regions, the host-responsive niche showed significantly increased proportions of fiber tracts (FB), hindbrain (HB), and the midbrain–hindbrain boundary (MB_HB), and was mainly enriched for monocytes and vascular-associated cell types (e.g., endothelial cells) (Figures S4B and S4C). This suggests that immune cells may migrate through fiber tracts and vascular regions. Spatial colocalization analysis revealed that monocytes showed stronger colocalization with vascular-associated cells, including endothelial cells and fibroblasts, within host-responsive niches (Figure S4D). Spatial gradient analysis further showed that monocytes were predominantly distributed near the center of the host-responsive niche, whereas they were mainly localized toward the boundary of the pathogen-infected niche (Figure S4E), suggesting distinct functional roles of the two niche types. Moreover, host-responsive niches exhibited increased expression of antiviral and inflammatory genes (e.g., *Ifng*, *Ccr1*, *S100a8*) (Figure S4F), and the upregulated genes were significantly enriched in immune response and chemotaxis-related processes (Figure S4G). Cell–cell communication analysis further revealed that, in host-responsive niches, monocytes showed strong communication with dendritic cells (Figure S4H). Overall, host-responsive niches are inflammatory response regions driven by vascular-mediated immune recruitment.

### STID reveals contrasting spatial niche organization in comparative analysis

To systematically characterize the structural and functional differences and spatiotemporal dynamics across distinct infection stages, STID established a multi-sample analytical framework supporting comparative and temporal analyses at the global (sample) and local (niche) levels (Figure 5A).

**Figure 5.**
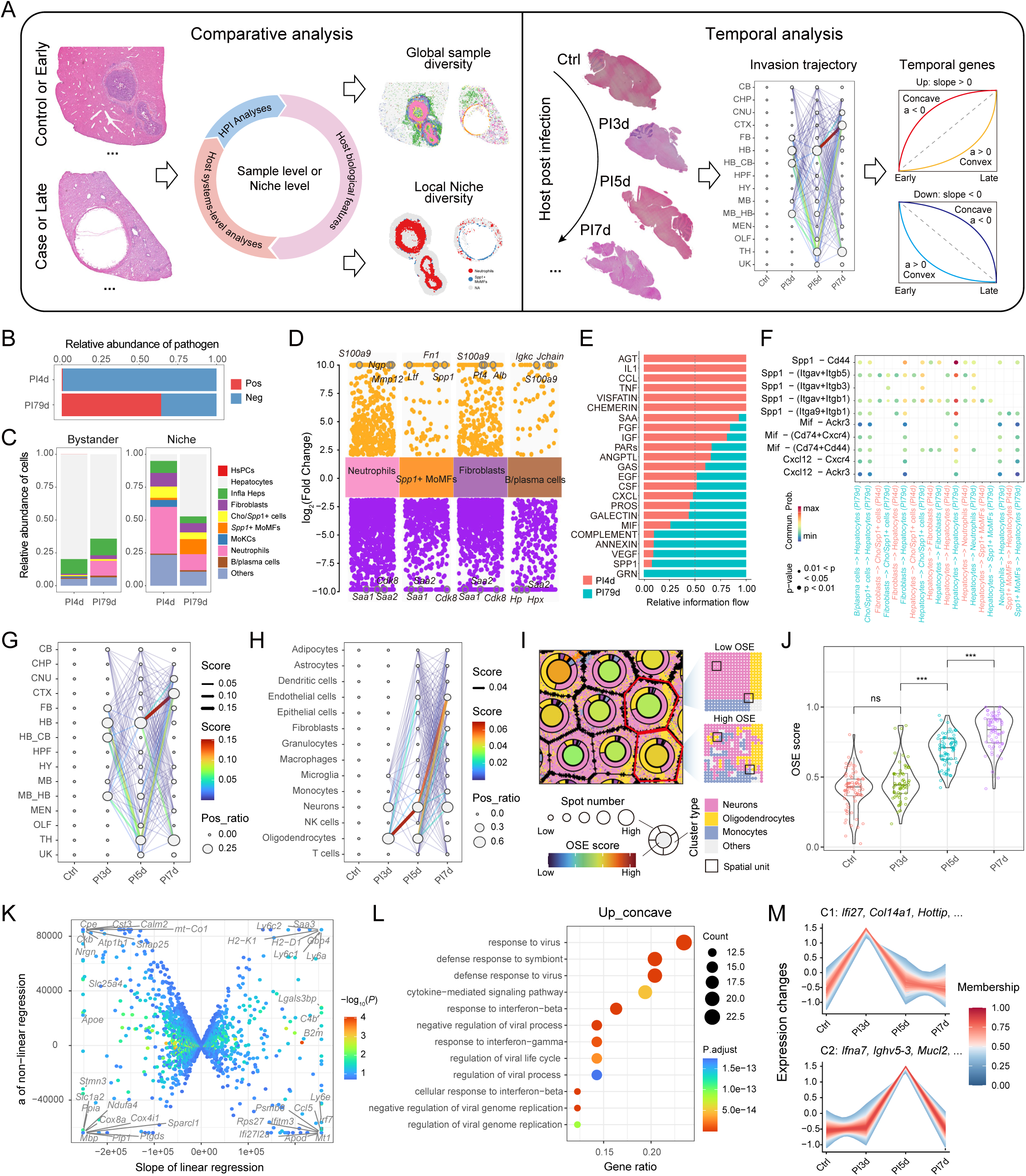
Infection-associated multi-sample comparative and temporal analyses. (A) Schematic overview of two representative study designs for multi-sample spatial transcriptomic analysis of infectious diseases. The left panel illustrates comparative analysis, used to compare control vs. disease or early vs. late stages. Comparative analysis is performed at both the sample level and niche level, integrating Host biological features, Host systems-level analyses, and HPI analysis to characterize global sample diversity and local niche diversity. The right panel illustrates temporal analysis, which is used to analyze consecutive infection time points. Temporal analysis is applied to characterize spatial invasion trajectories and temporally coordinated gene modules during infection progression. (B) Bar plot showing the proportion of pathogen-infected positive spots between PI4d (early) and PI79d (late) samples in the AE infection model. (C) Stacked bar plots showing differences in cellular composition between PI4d and PI79d samples in niche and bystander regions. Left: bystander region; right: niche region. (D) Volcano plot showing cell type-specific differentially expressed genes between PI4d and PI79d samples. (E) Bar plot showing relative information flow of major signaling pathways in cell–cell communication between PI4d and PI79d samples. (F) Bubble plot showing significantly altered ligand–receptor interaction pairs between PI4d and PI79d samples. (G–H) Bubble plots showing pathogen propagation patterns across tissues (G) and cell types (H) in the JE infection model. The y-axis represents tissues or cell types, and the x-axis represents post-infection time points. Bubble size indicates the proportion of pathogen-positive spots distributed across different tissues or cell types at each time point. Arrows indicate the inferred propagation paths. Arrow thickness and color intensity are proportional to the propagation score. Tissue abbreviations are defined as follows: TH, thalamus; CTX, cerebral cortex; CNU, cerebral nuclei; HB, hindbrain; FB, fiber tracts; CHP, choroid plexus; MB, midbrain; HY, hypothalamus; CB, cerebellum; HPY, hippocampal formation; MEN, meninges; OLF, olfactory bulb; MB_HB, midbrain–hindbrain boundary region; HB_CB, hindbrain–cerebellum boundary region; UK, unknown tissue. (I) Schematic overview of the organizational structure entropy (OSE) analysis workflow. (J) Box plots showing differences in OSE scores among Ctrl, PI3d, PI5d, and PI7d samples. Differences were assessed using the Wilcoxon rank-sum test. (K) Scatter plot showing trend classification of genes based on linear and nonlinear regression analysis. The x-axis represents the linear regression slope, and the y-axis represents the quadratic coefficient (a). Genes were classified into four trend categories based on their trajectory patterns: up-concave, up-convex, down-concave, and down-convex. (L) Bubble plot showing GO terms enriched among the top 100 genes ranked by slope within the up-concave category. (M) Line plots showing PI3d- and PI5d-specific gene modules identified by Mfuzz clustering, with plot titles indicating three representative genes with high membership scores. Top: PI3d; bottom: PI5d.

In the AE model, we applied STID to compare early (PI4d) and late (PI79d) stages. As disease progressed, the proportion of pathogen-infected spots significantly increased, accompanied by spatial remodeling of the microenvironment (Figures 5B and 5C). The early niche exhibited neutrophil-dominant acute inflammatory responses, whereas the late niche was enriched for *Spp1*□ MoMFs, indicating a gradual transition of the spatial microenvironment from acute inflammation to chronic inflammation, tissue repair, and remodeling. Meanwhile, neutrophils spread from the early niche into bystander regions at the late stage, indicating dissemination of inflammatory responses from localized niches to broader tissue regions (Figures 5C, S5A and S5B). In the late niche, the aggregation of *Spp1*□ MoMFs was markedly enhanced, whereas neutrophil aggregation was substantially reduced (Figure S5C). Spatial colocalization analysis further revealed that neutrophils were predominantly distributed around *Spp1*□ MoMFs, forming a hierarchical spatial organization centered on reparative cell populations (Figures S3A and S5D).

At the molecular level, differential expression analysis between late-stage and early-stage niches further supported this transition. In the late stage, acute inflammation-related genes (e.g., *Saa1* and *Saa2*) were globally downregulated, whereas *Spp1* expression in *Spp1*□ MoMFs was markedly increased, accompanied by significant upregulation of B cell–related genes (e.g., *Igkc* and *Jchain*) (Figures 5D and S5E). Functional enrichment analysis demonstrated that upregulated genes were primarily involved in tissue repair and remodeling, homeostatic maintenance, and humoral immunity, whereas downregulated genes were mainly enriched in metabolic pathways (Figure S5F). Cell–cell communication revealed a rewiring of signaling networks during disease progression (Figure 5E). The early niche was dominated by inflammatory signals such as IL1, CCL, TNF, and SAA, whereas the late niche shifted toward regulatory signals involved in tissue repair and remodeling, including SPP1, VEGF, and MIF. Ligand–receptor analysis further demonstrated that, in the late niche, the Spp1–(Itga/Itgb) signaling axis regulated extensive cell–cell interactions, primarily involving *Spp1*□ MoMFs, fibroblasts, and hepatocytes (Figure 5F). These findings suggest that SPP1-related signaling is predominantly driven by *Spp1*□ MoMFs and coordinates fibroblast activation and hepatocyte responses, thereby contributing to immune microenvironment remodeling and tissue repair processes.

### STID reveals spatiotemporal dynamics of pathogen infection in temporal analysis

In the JE infection model, we used longitudinal data to characterize the spatiotemporal dynamics of disease progression. By integrating samples across different stages, STID delineated temporal patterns of pathogen propagation, spatial tissue remodeling, and dynamic host transcriptional responses (Figure 5A).

As disease progressed, the proportion of pathogen-infected spots gradually increased, with hindbrain (HB) and thalamus (TH) showing the highest tissue abundance and oligodendrocytes and neurons constituting the dominant cell populations (Figures S5G and S5H). Pathogen propagation networks further revealed potential dissemination trajectories of Japanese encephalitis virus (JEV). At the tissue level, JEV was inferred to spread from the hindbrain and adjacent regions through the thalamus and progressively extend toward the cerebral cortex. At the cellular level, propagation trajectories primarily followed an oligodendrocyte-to-neuron-to-endothelial-cell axis (Figures 5G and 5H).

To characterize dynamic host responses during infection, we identified pathogen-infected and host-responsive niches across Ctrl, PI3d, PI5d, and PI7d samples (Figures S4A and S5I). Cellular composition analysis showed an increased proportion of endothelial cells, epithelial cells, fibroblasts, and monocytes in host-responsive niches rather than in pathogen-infected niches (Figures S5I–S5K). Infection also caused substantial microenvironmental remodeling, including immune cell infiltration, vascular remodeling, and fibrosis, predominantly within host-responsive niches. Moreover, organizational structure entropy (OSE) scores changed significantly as disease progressed, indicating continuous remodeling of spatial brain organization (Figures 5I, 5J, and S5L).

We next examined dynamic host transcriptional responses during infection progression. By jointly modeling linear and nonlinear temporal patterns, we classified genes into four categories: up-concave, up-convex, down-concave, and down-convex (Figures 5A and 5K). Functional enrichment analysis of the top 100 genes revealed that up-concave genes were primarily enriched in antiviral processes (e.g., response to virus), whereas up-convex genes were predominantly associated with antigen processing and presentation (Figures 5L and S5M). In contrast, down-convex genes were enriched in synaptic signaling and ion transport–related processes, whereas down-concave genes were associated with cytoskeletal homeostasis and energy metabolism. These results reveal temporally coordinated transcriptional programs underlying host responses during infection.

We further applied Mfuzz to identify stage-specific dynamic modules during intermediate infection stages. The PI3d-specific module included representative genes (e.g., *Ifi27*, *Col14a1*, and *Hoxa10*) associated with transient host remodeling during early infection. In contrast, the PI5d-specific module included representative genes (e.g., *Ifna7*, *Ighv5-3*, and *Mucl2*), consistent with a transition from early immune activation toward adaptive immunity and tissue remodeling (Figure 5M).

## Discussion

Systematically characterizing the structural and functional features of infection-associated spatial microenvironments is essential for understanding host–pathogen interactions that drive infection initiation, progression, and immune responses^9,12^. Here, we present STID, a standardized spatial transcriptomic analysis framework designed for infectious diseases. By integrating an infection-specific data structure with modular analytical workflows, including pathogen background correction, infection-associated spot detection, infection-associated niche identification, single-sample niche analysis, and multi-sample comparative and temporal analyses, STID provides a solution for standardized analysis of infectious microenvironments.

In infectious disease tissues, spatial transcriptomic data are often confounded by pathogen-derived background noise from sequencing contamination, background diffusion, and multi-mapped reads^22–24^. Such noise can obscure genuine infection signals, particularly in samples with low pathogen abundance or diffuse infection foci. STID implements a background correction strategy constrained by the distribution of pathogen gene expression in control samples. In both AE and TB models, this strategy effectively reduced nonspecific signal interference and enabled accurate detection of infection-associated spots based on the corrected expression profiles.

Different pathogen types and tissue contexts can generate pronounced spatial heterogeneity in infection-associated niches. To enhance the generalizability and robustness of STID, we further introduced a multi-mode niche identification strategy based on three spatial distribution patterns (Foci, Aggregated, and Dispersed), enabling the analysis framework to adapt to diverse infection-associated spatial architectures. Combined with density constraints and boundary expansion, STID supports refined delineation of niche boundaries. Across multiple infection models, including AE and JE, STID consistently identified pathogen-infected and host-responsive niches and revealed their spatial and functional segregation. Our results suggested that infectious microenvironments are spatially organized into distinct functional compartments, including pathogen-occupied regions, immune recruitment regions and immune response regions.

Beyond niche identification, STID further enhances multiscale characterization of infection-associated niches by integrating multiple layers of information, including tissue architecture, cellular composition, gene expression, cell–cell communication, and gene regulatory networks. In AE and JE models, STID revealed pronounced spatial stratification within niches, where distinct immune cell populations occupied specific spatial localizations, accompanied by structural and functional gradients radiating from the niche center to boundary. Furthermore, cell–cell communication and transcriptional regulatory networks formed coupled regulatory axes that jointly drove local inflammatory activation as well as tissue repair and remodeling processes. These findings uncover coordinated regulatory mechanisms underlying the functional compartmentalization of infection-associated niches.

As a comprehensive and robust analytical framework, STID also supports multi-sample comparative and temporal analyses, enabling systematic dissection of spatiotemporal dynamics during infection progression. Comparative analysis of early-and late-stage AE samples revealed a progressive transition from acute inflammation to chronic infection, accompanied by tissue repair and remodeling. This shift was accompanied by coordinated rewiring of immune cell composition, spatial organization, and cell–cell communication networks, reflecting the establishment of a pathologically adapted microenvironment. Temporal analysis of JE samples enabled extraction of temporal gene modules and reconstruction of pathogen dissemination trajectories, providing mechanistic insights into how infection remodels tissue architecture over time. These findings revealed the substantial structural and functional differences across different infection stages, which may correspond to stage-specific intervention strategies and guide the optimization of existing interventions.

Overall, by integrating a specialized data structure with the modular analytical framework, STID establishes a standardized, end-to-end workflow spanning from raw spatial transcriptomic data input to infection-associated single-sample and multi-sample analyses. Its broad applicability and robustness have been validated in multiple pathogen infection models. As spatial multi-omics technologies continue to be developed, STID is poised for extension toward multimodal data integration and modeling, thereby offering a computational foundation for investigations in pathogen biology, immunology, and translational medicine.

### Limitations of the study

Despite its utility, STID has several limitations. First, this study primarily relied on publicly available datasets, some of which lack high-quality cell-type annotations, limiting the depth of single-sample niche characterization and multi-sample analyses. Second, the delineation of pathogen-infected and host-responsive niches currently depends on gene expression and functional scoring. Therefore, further validation and refinement in larger, infection-associated spatial transcriptomic datasets are needed. Third, although parallel computing and sampling strategies improve computational efficiency, STID remains computationally demanding when applied to ultra-large spatial datasets (>100,000 spots). Fourth, STID offers limited support for imaging-based spatial transcriptomic data types, and cross-platform compatibility remains to be strengthened. Additionally, capturing the complexity of mixed infection states remains a challenge for current spatial transcriptomic approaches.

## Resource availability

### Lead contact

Further information and resource requests should be directed to Xiaofeng Shi.

### Materials availability

This study did not generate new unique reagents.

### Data and code availability

- All raw datasets used in this study were obtained from the original publications and are available through the Key Resources Table or Table S1. Processed data generated in this study are available in the Figshare repository (https://doi.org/10.6084/m9.figshare.31839988).
- The STID R package is publicly available on GitHub (https://github.com/YulongQin/STID) under the MIT License. The version used in this study has been archived on Zenodo (DOI: 10.5281/zenodo.20353070).
- All analysis scripts required to reproduce the results are available at: https://github.com/YulongQin/STID/tree/main/article_code.
- Documentation and user guidance for STID are available at: https://yulongqin.github.io/STID/.
- Any additional information required to reanalyze the data reported in this study is available from the lead contact upon reasonable request.

## Acknowledgments

This study was supported by the National Natural Science Foundation of China (Grant Nos. 82300527 and 82341118). The authors thank the International Symposium on Spatiotemporal Omics for Infectious Diseases (ISSOID) and all members of our laboratory for their support.

## Author contributions

Conceptualization, Y.Q. and X.S.; methodology, software, formal analysis, and data curation, Y.Q.; data curation, Y.P.; resources, Q.C., J.C., P.R., and Z.O.; writing – original draft, Y.Q.; writing – review & editing, Y.Q., Y.P., Q.C., H.D., D.W., X.L., and X.S.; supervision, Z.D. and X.S.

## Methods

### Key resources table

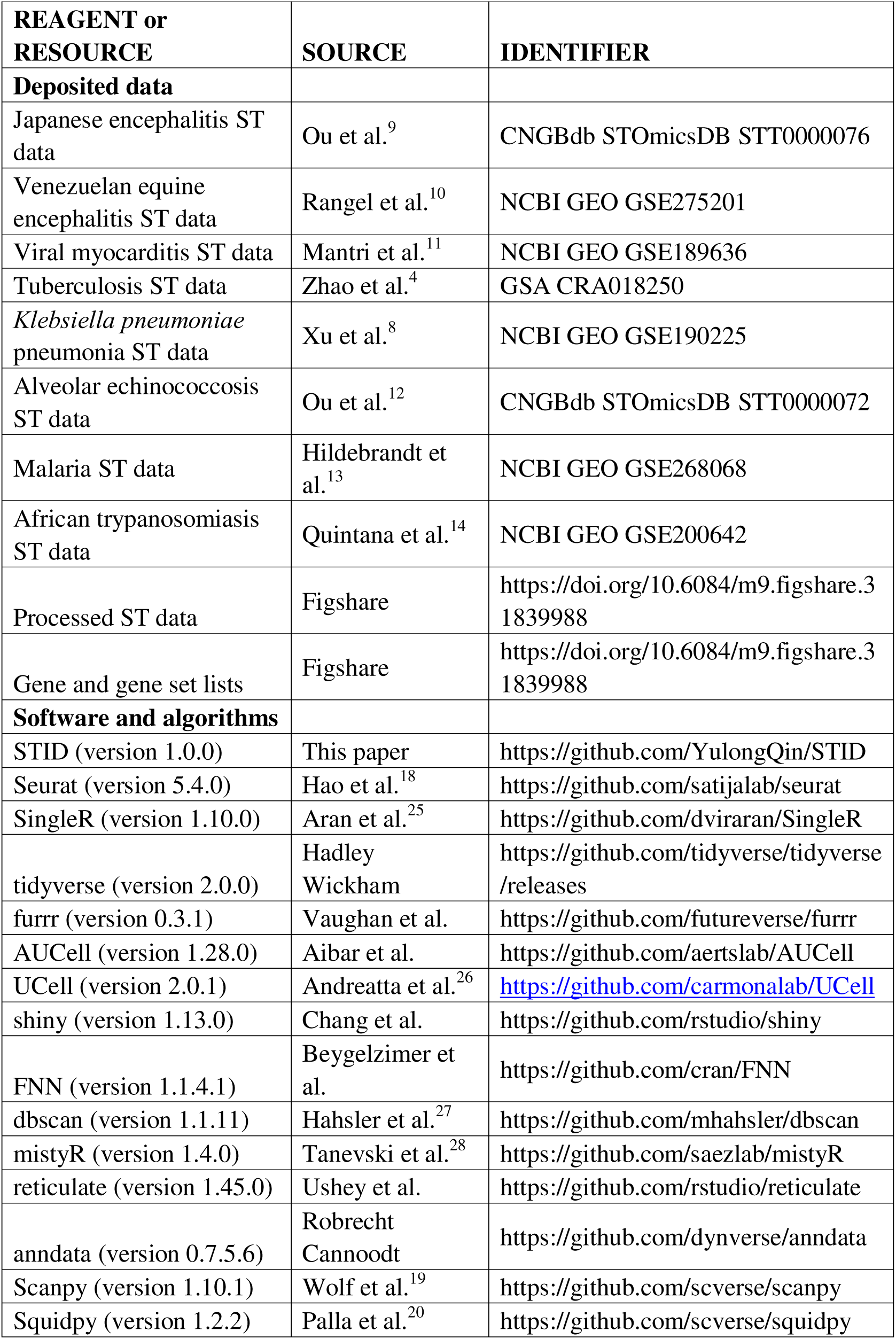
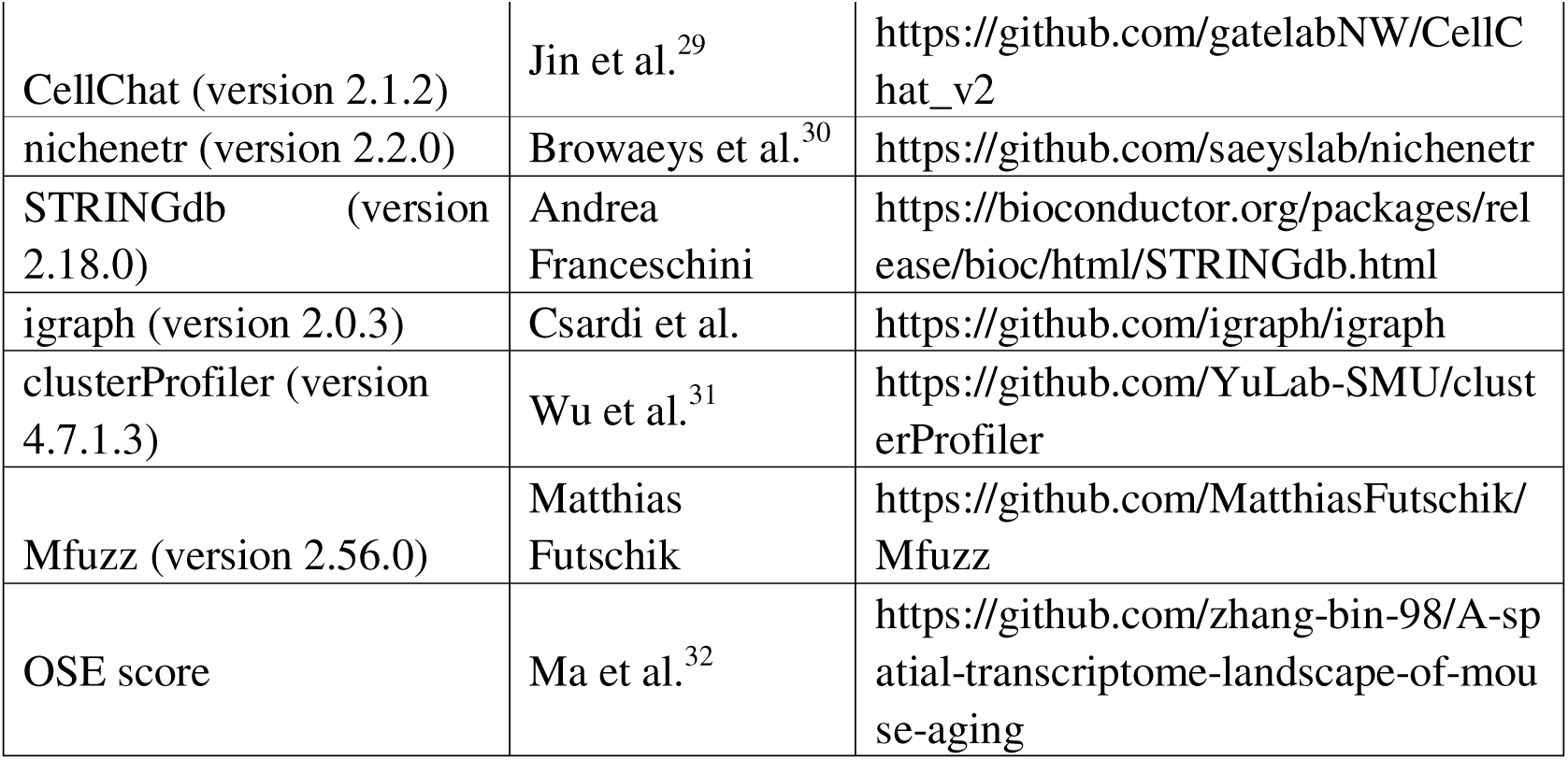

| **REAGENT or RESOURCE** | **SOURCE** | **IDENTIFIER** |
| --- | --- | --- |
| **Deposited data** |  |  |
| Japanese encephalitis ST data | Ou et al.<sup>9</sup> | CNGBdb STOmicsDB STT0000076 |
| Venezuelan equine encephalitis ST data | Rangel et al.<sup>10</sup> | NCBI GEO GSE275201 |
| Viral myocarditis ST data | Mantri et al.<sup>11</sup> | NCBI GEO GSE189636 |
| Tuberculosis ST data | Zhao et al.<sup>4</sup> | GSA CRA018250 |
| *Klebsiella pneumoniae* pneumonia ST data | Xu et al.<sup>8</sup> | NCBI GEO GSE190225 |
| Alveolar echinococcosis ST data | Ou et al.<sup>12</sup> | CNGBdb STOmicsDB STT0000072 |
| Malaria ST data | Hildebrandt et al.<sup>13</sup> | NCBI GEO GSE268068 |
| African trypanosomiasis ST data | Quintana et al.<sup>14</sup> | NCBI GEO GSE200642 |
| Processed ST data | Figshare | https://doi.org/10.6084/m9.figshare.31839988 |
| Gene and gene set lists | Figshare | https://doi.org/10.6084/m9.figshare.31839988 |
| **Software and algorithms** |  |  |
| STID (version 1.0.0) | This paper | https://github.com/YulongQin/STID |
| Seurat (version 5.4.0) | Hao et al.<sup>18</sup> | https://github.com/satijalab/seurat |
| SingleR (version 1.10.0) | Aran et al.<sup>25</sup> | https://github.com/dviraran/SingleR |
| tidyverse (version 2.0.0) | Hadley Wickham | https://github.com/tidyverse/tidyverse/releases |
| furrr (version 0.3.1) | Vaughan et al. | https://github.com/futureverse/furrr |
| AUCell (version 1.28.0) | Aibar et al. | https://github.com/aertslab/AUCell |
| UCell (version 2.0.1) | Andreatta et al.<sup>26</sup> | https://github.com/carmonalab/UCell |
| shiny (version 1.13.0) | Chang et al. | https://github.com/rstudio/shiny |
| FNN (version 1.1.4.1) | Beygelzimer et al. | https://github.com/cran/FNN |
| dbscan (version 1.1.11) | Hahsler et al.<sup>27</sup> | https://github.com/mhahsler/dbscan |
| mistyR (version 1.4.0) | Tanevski et al.<sup>28</sup> | https://github.com/saezlab/mistyR |
| reticulate (version 1.45.0) | Ushey et al. | https://github.com/rstudio/reticulate |
| anndata (version 0.7.5.6) | Robrecht Cannoodt | https://github.com/dynverse/anndata |
| Scanpy (version 1.10.1) | Wolf et al.<sup>19</sup> | https://github.com/scverse/scanpy |
| Squidpy (version 1.2.2) | Palla et al.<sup>20</sup> | https://github.com/scverse/squidpy |
| CellChat (version 2.1.2) | Jin et al.<sup>29</sup> | https://github.com/gatelabNW/CellChat_v2 |
| nichenetr (version 2.2.0) | Browaeys et al.<sup>30</sup> | https://github.com/saeyslab/nichenetr |
| STRINGdb (version 2.18.0) | Andrea Franceschini | https://bioconductor.org/packages/release/bioc/html/STRINGdb.html |
| igraph (version 2.0.3) | Csardi et al. | https://github.com/igraph/igraph |
| clusterProfiler (version 4.7.1.3) | Wu et al.<sup>31</sup> | https://github.com/YuLab-SMU/clusterProfiler |
| Mfuzz (version 2.56.0) | Matthias Futschik | https://github.com/MatthiasFutschik/Mfuzz |
| OSE score | Ma et al.<sup>32</sup> | https://github.com/zhang-bin-98/A-spatial-transcriptome-landscape-of-mouse-aging |

## Experimental model and study participant details

### External datasets

To evaluate the performance of STID modules, we collected and curated publicly available spatial transcriptomic datasets of infectious diseases, encompassing viral, bacterial, and parasitic infection models (Table S1). These datasets were generated from mouse brain, liver, heart, and lung tissues using Stereo-seq or 10x Visium platforms. Dataset-level metadata are summarized in Table S1.

## Method details

### Curation of gene and gene set resources

To support infection-associated spatial transcriptomic analysis, we curated pathogen infection-related and host immunity-related gene and gene set resources and integrated them into the STID R package (Table S2). Gene lists included immune response genes and genes related to pattern recognition receptors (PRRs), cytokines, and chemokines. Human and mouse gene annotations were obtained from GENCODE v46 and GENCODE vM35, respectively. Gene set lists were compiled from GO Biological Process, KEGG, Reactome, and MSigDB Hallmark gene sets, with additional inclusion of programmed cell death (PCD)-related pathways. Corresponding human and mouse ortholog annotations were provided. For infection models lacking matched disease-specific pathways in public databases, biologically related disease signatures, such as malaria- or Chagas disease-related signatures, were used as surrogate host-response signatures.

### Pathogen gene background correction

To reduce background noise from pathogen genes in spatial transcriptomic data, we applied a control-based background correction strategy. For each pathogen gene, the 95th percentile of nonzero expression values in background samples was calculated and used as the background expression threshold. To account for differences in sequencing depth, this threshold was further adjusted using the ratio of average unique molecular identifier (UMI) counts between background samples and all samples. The adjusted threshold was then subtracted from gene expression values across all spots, with negative values set to zero.

### Infection-associated spot detection

The detection of infection-associated spots was performed using two complementary strategies: a gene expression–driven approach and a gene set score–driven approach. For the gene expression–driven approach, the expression distribution of pathogen infection- or host response–related genes was calculated within each sample, and the 5th percentile of non-zero expression values was used as the default threshold. This threshold could be further adjusted based on the spatial distribution of gene expression and inflection points in frequency histograms. Spatial spots with expression values exceeding the threshold were defined as infection-associated positive spots. For multiple genes, aggregated expression across genes was used for detection. For the gene set score–driven approach, gene set scores were computed using algorithm-based methods (AddModuleScore, AUCell, and UCell) or expression-based metrics (mean or total gene expression), and thresholds were defined using the same strategy as in the gene expression–driven approach. Spots with scores exceeding the threshold were designated as positive spots. Both strategies support optional data smoothing prior to threshold determination to reduce local noise.

### Infection-associated niche identification

To systematically identify infection-associated niches in spatial transcriptomic data, we constructed a unified niche detection framework based on the spatial aggregation patterns of positive spots. STID provides three niche identification modes—Foci, Aggregated, and Dispersed—corresponding to high, intermediate, and low levels of spatial aggregation, respectively.

For Foci-type niches, the presence of visually distinguishable infectious foci with clear boundaries was first confirmed using H&E staining. After histological confirmation, *NicheDetect_Lasso* was applied for interactive delineation of foci-associated niches. This method integrates multiple auxiliary features, including pathogen-infected spots, host-responsive spots, and spatial distributions of cell type annotations, together with H&E histology as a gold standard, to accurately define infection-associated niches.

For Aggregated-type niches, automated identification was performed using the *NicheDetect_STS* function. Spatial transcriptomic data were first preprocessed, including low-density region filtering based on kernel density estimation and iterative refinement of positive spots via k-nearest neighbor (kNN)–based neighborhood propagation, to reduce background noise and enhance local spatial signals. On this basis, either region-level or spot-level strategies were applied for niche identification according to the degree of spatial aggregation among processed spots.

In region-level mode, DBSCAN density clustering was first performed on processed spots to identify putative spatial aggregates, with clustering parameters adaptively determined based on spot distribution characteristics. Cluster boundaries were then reconstructed using either convex hull or concave hull methods to define infection-associated niche regions. In spot-level mode, spatial adjacency graphs were constructed using kNN, followed by connected component analysis to identify spatially contiguous spot clusters, enabling precise delineation of infection-associated niches.

For Dispersed-type niches, *NicheDetect_Spot* was applied. Each positive spot was treated as an independent analytical unit, and all positive spots were directly defined as niches.

Additionally, *NicheExpand* was optionally applied to refine initial niches. Specifically, the distance from each bystander spot to the nearest ROI boundary was computed, and spots within a predefined distance threshold were incorporated into the corresponding niche ROIs, enabling flexible expansion of niche regions.

### Niche center and boundary determination

After niche identification, boundary and center spots were determined for each ROI. Boundary spots were identified based on k-nearest neighbor (kNN) relationships. For each spot within a niche, if the number of neighboring spots assigned to other niches was greater than or equal to a predefined threshold (k = 8 for square-grid data, k = 6 for hex-grid data, threshold = 1), the spot was labeled as a boundary spot. Center spots were defined based on the median x and y coordinates of all spots within each ROI. The spot closest to the coordinate median was selected as the center; if the distance exceeded 5× the coordinate spacing, the coordinate median was directly used as the center. Distances from all spots to the nearest center were then calculated, and bystander spots were assigned to the nearest ROI.

### Spatial gradient analysis

To characterize spatial patterns of variation from niche centers toward bystander regions and to identify potential spatial gradient transition points, STID provides two types of spatial gradient analyses: gene expression–based and cell composition–based approaches. Gene expression gradient analysis uses the spatial distance to the niche center as a continuous variable to describe changes in target genes or features along spatial gradients. Optionally, expression values can be Z-score normalized, and distances can be normalized using quantile-based scaling. Generalized additive models (GAMs) were then applied to smooth spatial gradients, and the fitted results were visualized across samples, with niche boundaries annotated. Cell composition gradient analysis was performed using distance-based binning. Spatial distances were adaptively partitioned into multiple bins according to spatial resolution, and the relative proportions of different cell types within each bin were calculated. The results were presented as fitted spatial gradient curves, with niche boundaries annotated.

### Cell aggregation index

To more accurately characterize cellular aggregation across spatial regions, STID refines a previously proposed cell aggregation index (CAI)^33^. For each cell type within a given region, we define a local aggregation measure based on the mean spatial distance between each cell and its k nearest same-type neighbors (k = 8 for square-grid data and k = 6 for hex-grid data). This distance is then transformed into a single-cell aggregation score using an exponential function, and the regional CAI is defined as the average of all single-cell scores.

Specifically, for each cell i, the local distance is defined as:

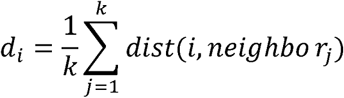

The single-cell aggregation index is defined as:

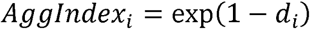

and the regional cell aggregation index is defined as:

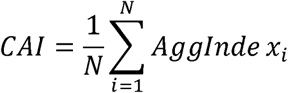

where i denotes the i-th cell (i = 1, 2, …, N); k denotes the number of nearest neighbors used for local aggregation estimation; dist(i, j) denotes the spatial distance between cell i and its j-th nearest same-type neighbor; N denotes the total number of cells of a given type within the region; d_i denotes the mean distance between cell i and its k nearest same-type neighbors; AggIndex_i denotes the single-cell aggregation score; and CAI denotes the mean aggregation index within the region.

To further quantify the spatial extent of aggregated cells, a cell adjacency graph is constructed, and spatially connected components are identified. Connected clusters containing at least two nodes are retained as valid aggregated groups. The proportion of cells contained in these groups relative to all cells of the same type is defined as the cell high-aggregation region proportion, which reflects the spatial enrichment level of each cell type.

### Colocalization analysis

To systematically assess spatial colocalization between cells or gene expression patterns, STID provides two complementary approaches: a spatial proximity–based method and an expression-based spatial modeling approach. The spatial proximity–based method uses Squidpy (via reticulate in R) for analysis. Neighborhood enrichment and co-occurrence are computed to quantify local co-occurrence patterns of cells or genes in spatial contexts. The expression-based method integrates mistyR for multiview modeling, constructing intra-view, juxtaview, and paraview spatial views to assess the contribution of spatial information at different scales to the prediction of cell- or gene-level features. The improvement in model explanatory power is quantified using ΔR² and ΔRMSE. In addition, STID provides two-dimensional colocalization visualizations for cell-type or gene pairs to illustrate their spatial distribution patterns.

### Differential gene expression and gene set enrichment analysis

Differential gene expression analysis was performed using Seurat to compare transcriptional differences between samples or spatial regions at both the global and cell-type levels. The Wilcoxon rank-sum test was used to identify differentially expressed genes, followed by Benjamini–Hochberg correction for multiple testing. Differential results were visualized using volcano plots at both the global and cell-type levels.

Gene set enrichment analysis was conducted using clusterProfiler, including over-representation analysis (ORA) and gene set enrichment analysis (GSEA). ORA was performed on differentially expressed genes or module genes to identify enriched Gene Ontology (GO) terms and KEGG pathways. GSEA was performed on a ranked list of all genes based on log2 fold change to assess pathway enrichment patterns across the expression profile. Enrichment results were visualized using bubble plots.

### Cell–cell communication and gene regulatory network analysis

Cell–cell communication analysis was performed using CellChat by integrating ligand–receptor expression information with spatial constraints to infer intercellular communication strengths within different samples or spatial regions. Weighted cell–cell interaction networks were constructed to identify key cell types, followed by analysis of major signaling pathways and ligand–receptor interactions among these cells.

Gene regulatory network analysis was performed using NicheNet to construct ligand-driven intercellular regulatory networks. Candidate ligands were prioritized by integrating receiver-cell transcriptomic profiles, sender-cell ligand expression, and prior ligand–receptor interactions, followed by prediction of their downstream target genes. Multilayer regulatory networks were then constructed by integrating ligand–receptor interactions, signal transduction networks, and ligand–regulator regulatory relationships, linking ligand inputs to regulator activity and downstream target gene expression. This framework enables the characterization of signal propagation from sender to receiver cells within niches at the level of transcriptional regulation.

### Host–pathogen interaction analysis

STID provides two complementary strategies to analyze host–pathogen interactions: correlation-based analysis at the transcriptomic level and protein interaction network–based inference. At the transcriptomic level, correlations between pathogen and host genes were computed using gene expression profiles derived from spatial niches or defined cell populations. Host genes significantly associated with pathogen genes were selected for downstream functional analysis.

At the protein level, FASTA sequences of pathogen and host proteins were retrieved, and BLAST (blastp) was used for sequence alignment. High-confidence homologous matches were identified based on sequence similarity metrics including E-value, percent identity, bit score, and alignment length, yielding candidate cross-species protein correspondences. Organism-specific STRING database files (protein alias, annotation, and interaction networks) were retrieved, and BLAST-matched proteins were mapped to STRING IDs to obtain known host protein–protein interactions, which were used to infer potential host–pathogen interaction links. A high-confidence protein–protein interaction network was then constructed using a threshold on the STRING combined score, from which host genes associated with pathogen proteins were extracted for downstream functional analysis.

### Pathogen invasion trajectory analysis

To characterize the spatiotemporal propagation of pathogens across different tissues or cell types during infection, STID constructed an infection propagation score matrix based on adjacent time points to infer potential pathogen invasion trajectories. For two consecutive time points (*t* and *t+1*), the proportional composition of each tissue or cell type within pathogen-infected spots at time point *t* was first calculated (*Load_source,t_*). Subsequently, the change in the proportion of the target tissue or cell type within pathogen-infected spots over the interval *t*→*t+1* was quantified as:

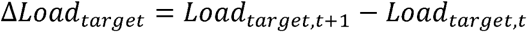

Based on these definitions, a propagation score was calculated as:

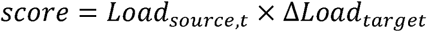

Only propagation events with Δ*Lood_target_* >0 were retained to focus on potential propagation events associated with infection expansion. Propagation scores across different time points and tissue or cell types were then integrated to construct a pathogen propagation network. In this network, arrow direction indicates the inferred direction of propagation, whereas edge width and color intensity represent the magnitude of the propagation score, enabling intuitive visualization of potential pathogen invasion trajectories during infection.

### Spatial organizational entropy analysis

To quantify dynamic changes in spatial structural disorder during infection, STID refined a previously proposed Organizational Structure Entropy (OSE) framework^32^. OSE quantifies spatial disorganization by measuring heterogeneity in local tissue organization and cellular composition.

Spatial samples were first segmented into superpixels using Simple Linear Iterative Clustering (SLIC), where each superpixel represents a region with spatially and transcriptionally coherent features. Within each superpixel, a local spatial unit was defined as a central spot together with its neighboring spots. To ensure robustness, local spatial units with fewer than three neighboring spots were excluded to reduce noise from edge or sparse regions.

The observed spatial entropy of each region was computed as:

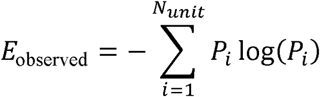

where *Pi* denotes the frequency of the i-th unit type, and *N*_unit_ represents the number of unit types observed in the region.

To account for differences in spot numbers across superpixels and ensure comparability of entropy values, a normalization factor based on the expected number of unit types was introduced. The expected number of unit types was defined as:

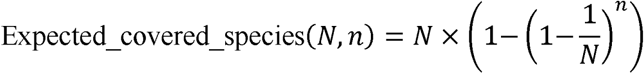

where *N* denotes the total number of possible unit types in the region, and *n* denotes the number of observed local spatial units.

The adjusted spatial entropy was then calculated as:

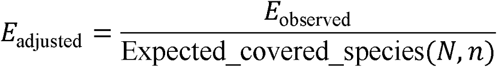

### Temporal gene module identification

To characterize the dynamic remodeling of host transcriptional responses during infection, STID developed a temporal gene module identification framework to identify dynamic transcriptional programs across multiple biological levels, including whole samples, infection-associated niches, and defined cell populations. Gene expression matrices were first grouped according to temporal labels, and average gene expression profiles at each time point were calculated to construct temporal expression matrices.

STID then provides two complementary strategies for identifying dynamic transcriptional programs. The first strategy is based on trend fitting, in which dynamic genes with significant temporal dependence are identified by combining linear regression and quadratic polynomial regression. Linear regression was used to evaluate the direction of expression changes, whereas quadratic polynomial models were used to characterize nonlinear dynamic trends. Based on the linear regression coefficient (slope) and the quadratic term coefficient (*a*), dynamic genes were classified into four expression patterns: up-concave (slope > 0 and *a* < 0), up-convex (slope > 0 and *a* > 0), down-concave (slope < 0 and *a* < 0), and down-convex (slope < 0 and *a* > 0).

The second strategy is based on fuzzy clustering (Mfuzz) and is designed to identify gene modules with complex temporal expression trajectories. Following missing value imputation, variance filtering, and normalization, fuzzy c-means clustering was performed to calculate gene–module membership scores, and core module genes were identified based on predefined thresholds.

### Quantification and statistical analysis

Statistical significance between groups was assessed using the Wilcoxon rank-sum test in R (v4.2.0). Data are presented as mean ± SEM unless otherwise indicated. Asterisks denote statistical significance as follows: \**P* < 0.05, \*\**P* < 0.01, and \*\*\**P* < 0.001.

---

**Figure S1.**
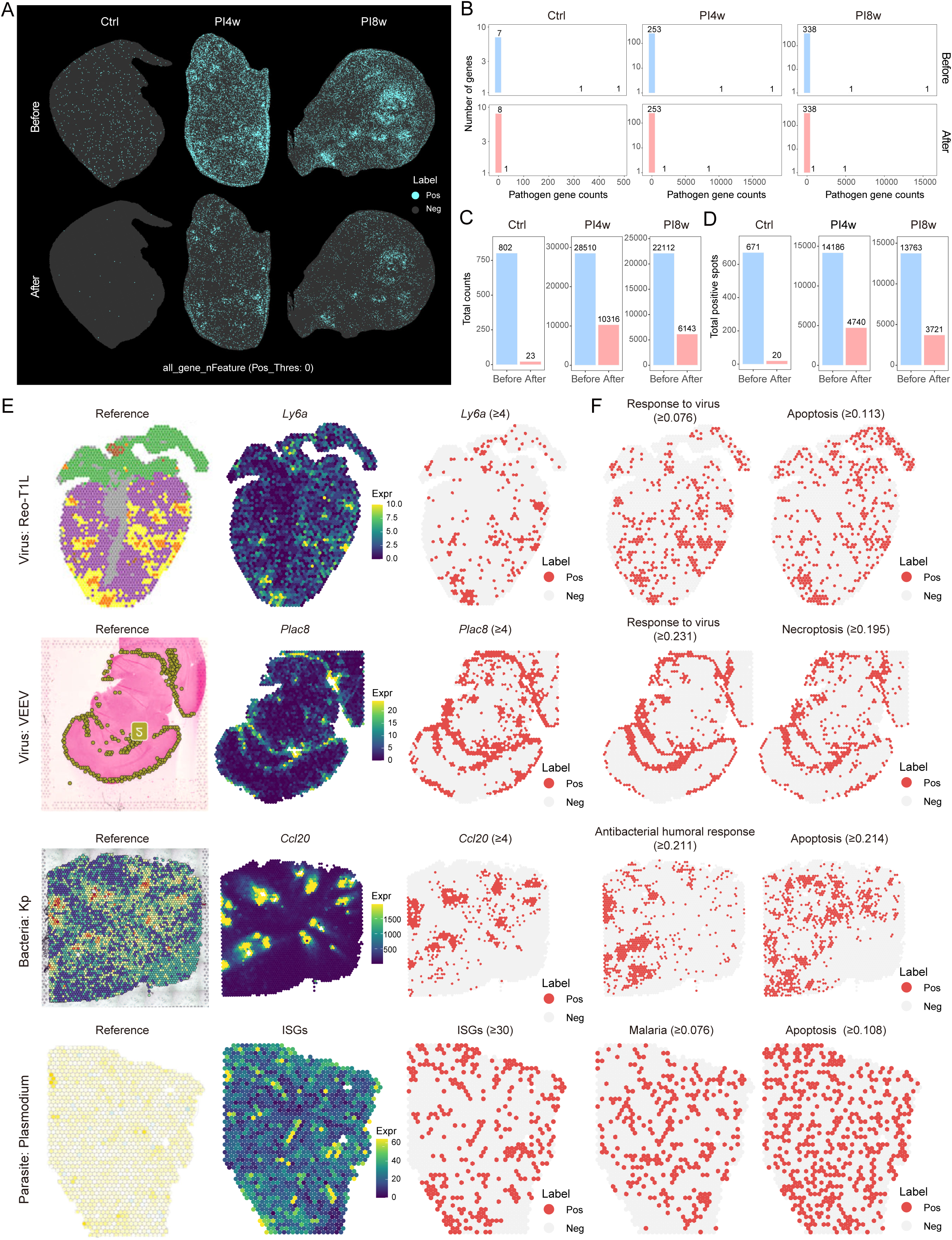
Validation of pathogen background correction and infection-associated spot detection. (A) Spatial maps showing pathogen gene expression in tuberculosis model samples before and after pathogen background correction (Control, PI4w, and PI8w stages). Top: before correction; bottom: after correction. (B) Histograms showing frequency distributions of total pathogen gene counts in samples at different infection stages. Top: before correction; bottom: after correction. (C–D) Bar plots showing changes in total pathogen gene counts and numbers of pathogen-infected spots across infection stages before and after pathogen background correction. (E) Spatial maps showing detection of host-responsive spots across different infection models (viral myocarditis, Venezuelan equine encephalitis, *Klebsiella pneumoniae* pneumonia, and malaria). Left: published results; middle: representative host-responsive gene expression; right: identified host-responsive spots. (F) Spatial maps showing other types of host-responsive spots across infection models, including anti-pathogen response score-positive spots and cell death score-positive spots.

**Figure S2.**
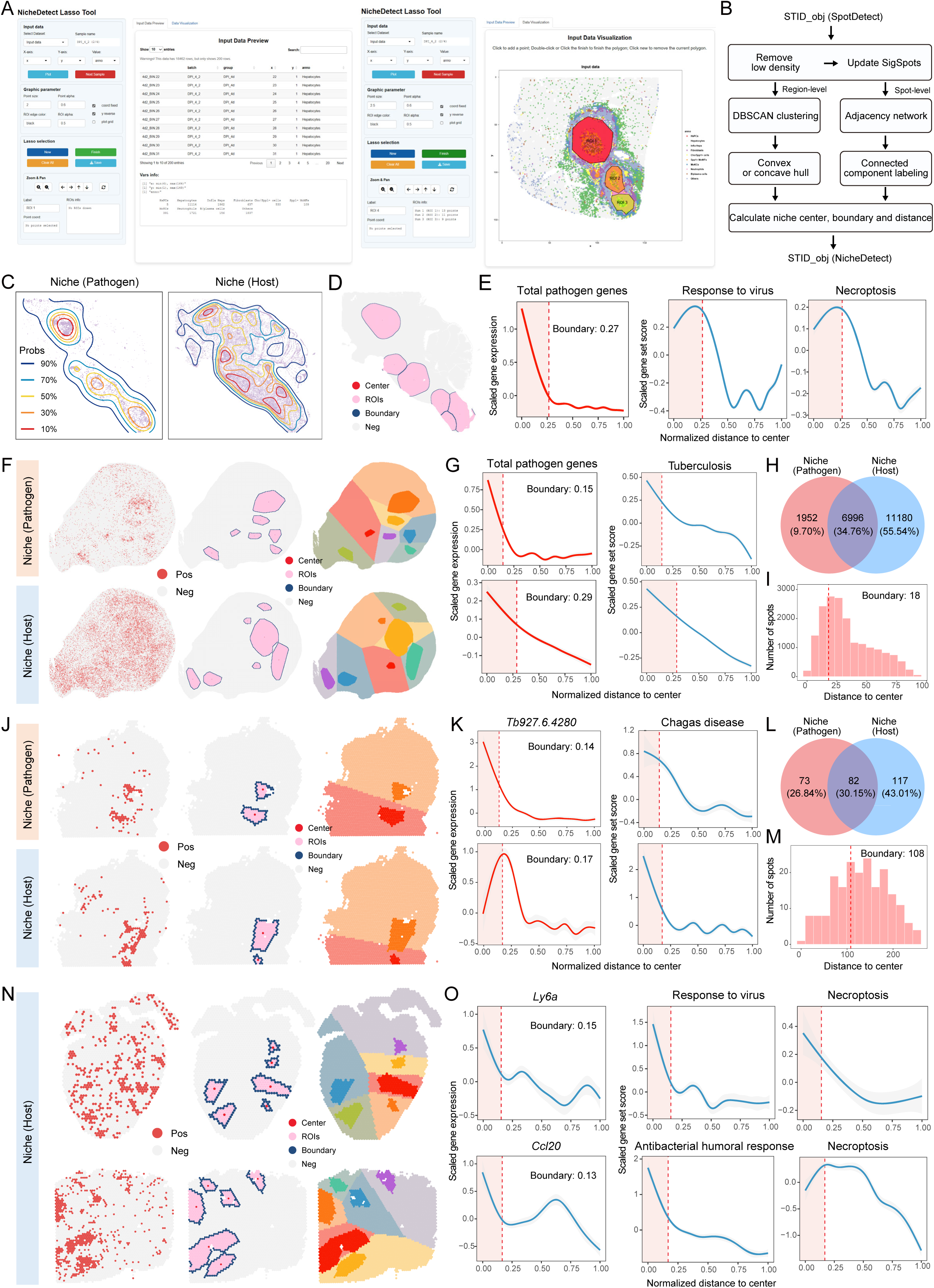
Validation of identification of infection-associated niches. (A) Shiny interface of NicheDetect_Lasso for interactive identification of niches in the AE model, including data visualization and lasso-based selection panels. (B) Schematic workflow of the NicheDetect_STS method. Key steps include removal of low-density spots and updating significant spots, followed by region-level (DBSCAN clustering and convex/concave hull) and spot-level analyses (adjacency network and connected components), and ROI-based center, boundary, and distance calculations. (C) Spatial maps showing the density of pathogen-infected and host-responsive spots after low-density filtering in the JE model. Left: pathogen; right: host. (D) Spatial maps showing niche structures after boundary expansion. (E) Fitted curves showing spatial gradient of total pathogen gene counts, response to virus, and necroptosis from niche centers toward the periphery after boundary expansion. (F) Spatial maps showing pathogen-infected niche and host-responsive niche in the tuberculosis model. Top: pathogen-infected niche; bottom: host-responsive niche. Left: positive spot distribution; middle: niche structures; right: multiple ROIs within the niche and corresponding bystander regions. (G) Fitted curves showing spatial gradient of total pathogen gene counts and tuberculosis score from niche centers toward the periphery. Dashed lines indicate niche boundaries. (H) Venn diagram showing spatial overlap between pathogen-infected and host-responsive niches. (I) Histogram showing the distribution of distances from host-responsive niche spots to the nearest pathogen-infected niche center. Dashed lines indicate niche boundaries. (J) Spatial maps showing pathogen-infected niche and host-responsive niche in the African trypanosomiasis model. Top: pathogen-infected niche; bottom: host-responsive niche. Left: positive spot distribution; middle: niche structures; right: multiple ROIs within the niche and corresponding bystander regions. (K) Fitted curves showing spatial gradient of total pathogen gene counts and Chagas disease scores from niche centers toward the periphery. Dashed lines indicate niche boundaries. (L) Venn diagram showing spatial overlap between pathogen-infected and host-responsive niches. (M) Histogram showing the distribution of distances from host-responsive niche spots to the nearest pathogen-infected niche centers. Dashed lines indicate niche boundaries. (N) Spatial maps showing host-responsive niches in viral myocarditis and *Klebsiella pneumoniae* pneumonia models. Top: viral myocarditis; bottom: *Klebsiella pneumoniae* pneumonia. Left: positive spot distribution; middle: niche structures; right: multiple ROIs within the niche and corresponding bystander regions. (O) Fitted curves showing spatial gradient of host response features from niche centers toward the periphery. Dashed lines indicate niche boundaries. Top: viral myocarditis; bottom: *Klebsiella pneumoniae* pneumonia.

**Figure S3.**
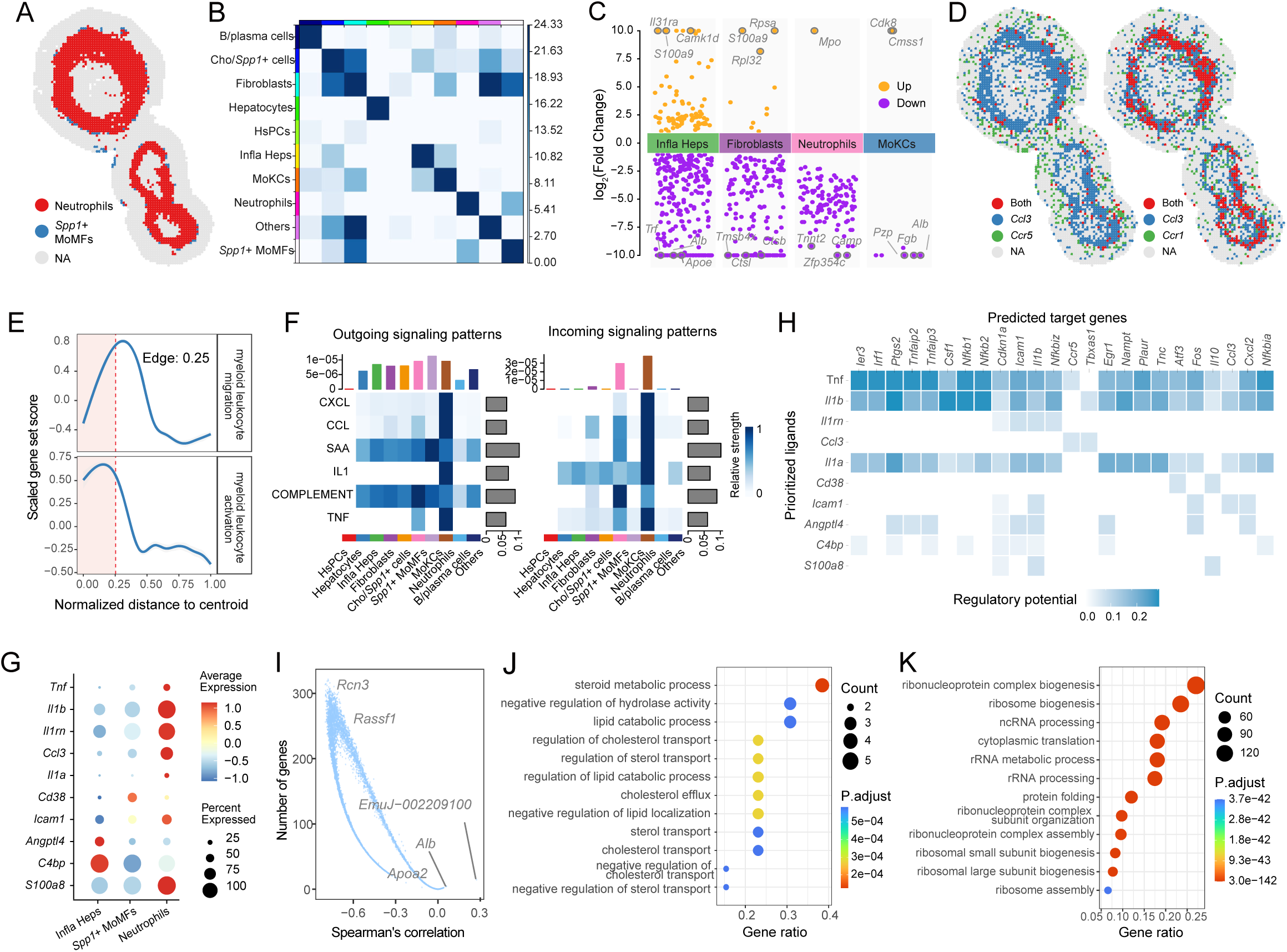
Supplementary multiscale single-sample niche analysis. (A) Spatial maps showing colocalization between neutrophils and *Spp1*□ MoMFs within the niche in the AE model. (B) Spatial maps showing colocalization of ligand–receptor pairs within the niche. Left: *Ccl3* – *Ccr1*; right: *Ccl3* – *Ccr5*. (C) Heatmap showing spatial colocalization between different cell types within the niche inferred using *Squidpy*. (D) Volcano plot showing differential gene expression analysis between the niche and bystander region across all cell types. (E) Fitted curves showing spatial gradients of GO-term gene-set scores enriched among differentially expressed genes from niche centers toward the periphery. Dashed lines indicate niche boundaries. Top: myeloid leukocyte migration; bottom: myeloid leukocyte activation. (F) Heatmap showing signaling strengths of major pathways across all cell types within the niche. Left: outgoing signaling patterns; right: incoming signaling patterns. (G) Bubble plot showing expression of key ligand genes across different cell types. (H) Heatmap showing ligand–receptor regulatory potential scores between sender cells and target cells. Sender cells include inflammatory hepatocytes, neutrophils, and *Spp1*□ MoMFs; target cells include neutrophils, *Spp1*□ MoMFs, MoKCs, and fibroblasts. (I) Dot plot showing Spearman correlations between host and pathogen gene expression. (J) Bubble plot showing GO terms enriched among host genes positively correlated with pathogen gene expression. (K) Bubble plot showing GO terms enriched among host genes potentially interacting with pathogen genes.

**Figure S4.**
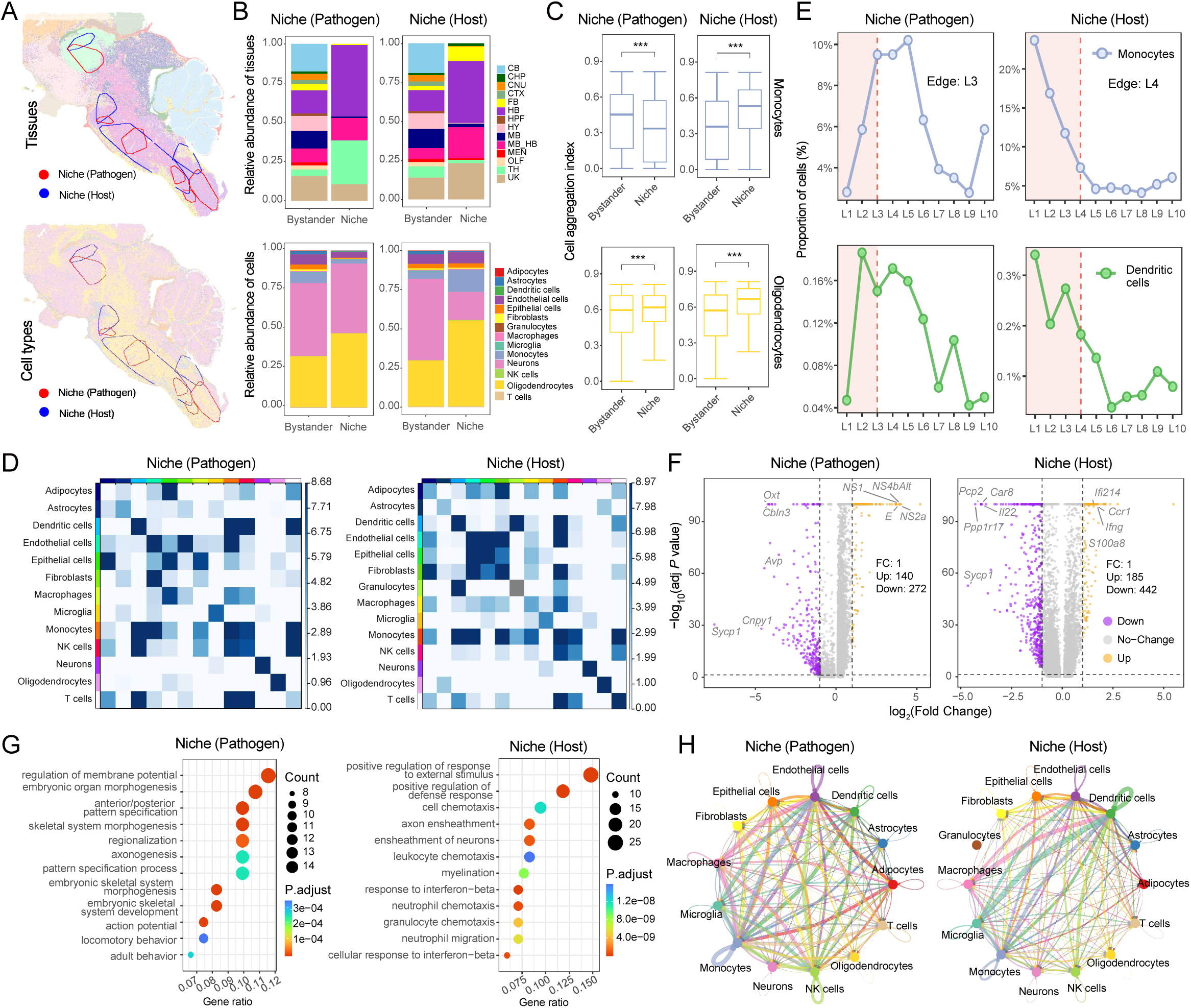
Comparison of pathogen-infected and host-responsive niches. (A) Spatial maps showing the distribution of tissue and cell types in the JE model, with pathogen-infected and host-responsive niches indicated in red and blue, respectively. Top: tissue types; bottom: cell types. Tissue abbreviations are defined as follows: TH, thalamus; CTX, cerebral cortex; CNU, cerebral nuclei; HB, hindbrain; FB, fiber tracts; CHP, choroid plexus; MB, midbrain; HY, hypothalamus; CB, cerebellum; HPY, hippocampal formation; MEN, meninges; OLF, olfactory bulb; MB_HB, midbrain–hindbrain boundary region; HB_CB, hindbrain–cerebellum boundary region; UK, unknown tissue. (B) Stacked bar plots showing the tissue and cellular composition of the pathogen-infected and host-responsive niches. Left: pathogen-infected niche; right: host-responsive niche. Top: tissue composition; bottom: cell composition. (C) Boxplots showing the cell aggregation index in the pathogen-infected and host-responsive niches. Top: monocytes; bottom: oligodendrocytes. (D) Heatmaps showing spatial colocalization among all cell types within the pathogen-infected and host-responsive niches. (E) Line plots showing spatial gradient of cell type proportions from niche centers toward the periphery in the pathogen-infected and host-responsive niches. Dashed lines indicate niche boundaries. (F) Volcano plots showing differentially expressed genes in the pathogen-infected and host-responsive niches relative to their respective bystander regions. (G) Bubble plots showing GO terms enriched among upregulated differentially expressed genes in the pathogen-infected and host-responsive niches. (H) Network diagrams showing cell–cell communication weights among all cell types within the pathogen-infected and host-responsive niches.

**Figure S5.**
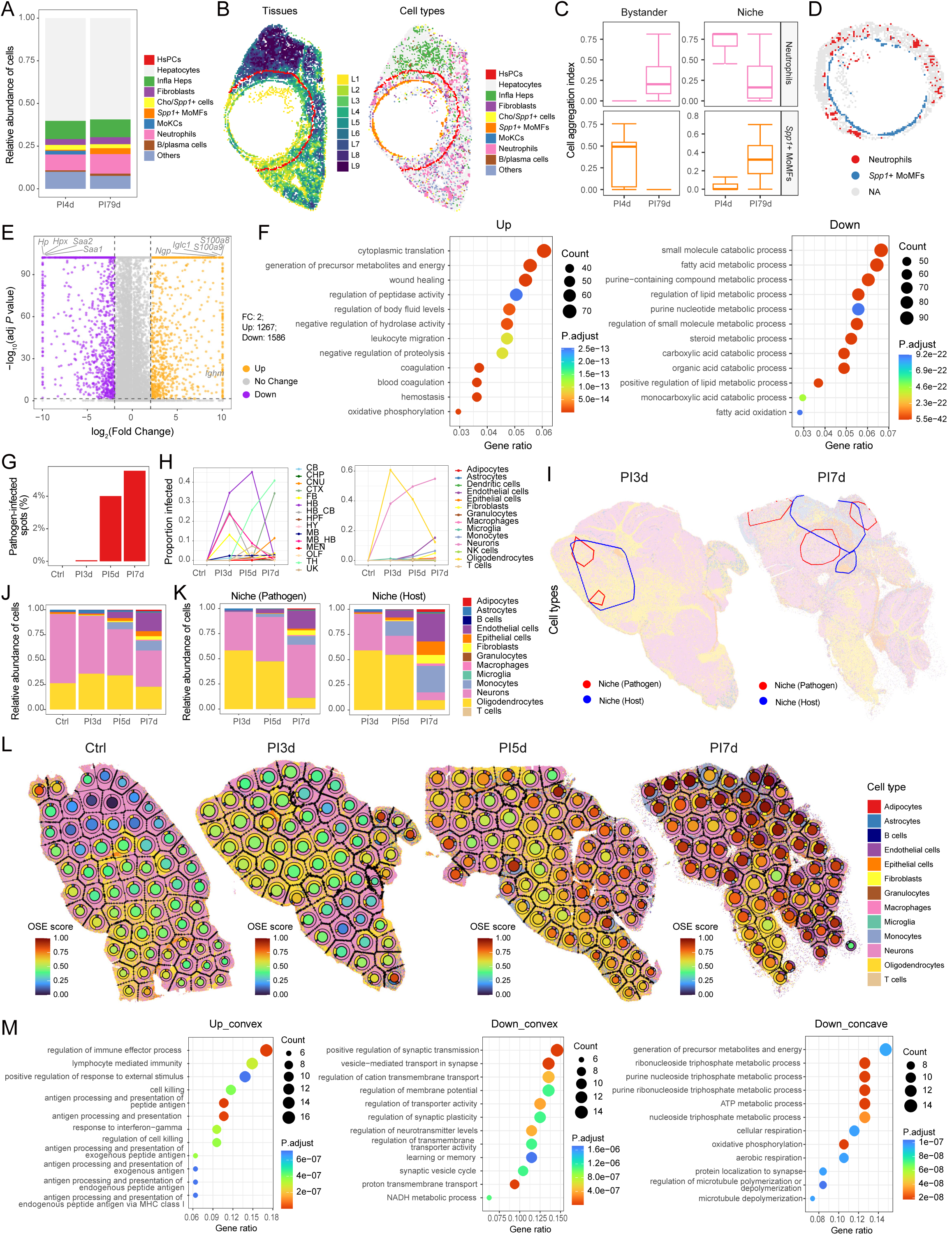
Supplementary infection-associated multi-sample comparative and temporal analyses. (A) Stacked bar plots showing differences in cellular composition between PI4d (early) and PI79d (late) samples in the AE infection model. (B) Spatial maps showing liver zonation and spatial distribution of cell types in PI79d samples. Red circles indicate niche boundaries. Left: liver zonation; right: cellular composition. (C) Boxplots showing differences in cell aggregation index between niche and bystander regions in PI4d and PI79d samples. Top: neutrophils; bottom: *Spp1*□ MoMFs. Left: bystander region; right: niche region. (D) Spatial maps showing spatial colocalization of neutrophils and *Spp1*□ MoMFs in PI79d samples. (E) Volcano plot showing differentially expressed genes between PI4d and PI79d samples. (F) Dot plot showing GO terms enriched among upregulated and downregulated differentially expressed genes. Left: upregulated genes; right: downregulated genes. (G) Bar plots showing the proportion of pathogen-infected spots across different infection time points. (H) Line plots showing changes in the proportions of tissue and cell types among pathogen-positive spots across different infection stages in the JE infection model. (I) Spatial maps showing the distribution of cell types within pathogen-infected and host-responsive niches in the JE infection model at PI3d and PI7d, with pathogen-infected niches and host-responsive niches indicated in red and blue, respectively. (J) Stacked bar plots showing the cellular composition of samples across different infection time points. (K) Stacked bar plots showing the cellular composition within pathogen-infected and host-responsive niches across different infection time points. Left: pathogen-infected niches; right: host-responsive niches. (L) Spatial maps showing OSE scores and cellular composition across different local spatial units in Ctrl, PI3d, PI5d, and PI7d samples. (M) Bubble plots showing GO terms enriched among the top 100 genes ranked by slope within the up-convex, down-convex, and down-concave categories.

